# Multiple and subject-specific roles of uncertainty in reward-guided decision-making

**DOI:** 10.1101/2024.03.27.587016

**Authors:** Alexander Paunov, Maëva L’Hôtellier, Dalin Guo, Zoe He, Angela Yu, Florent Meyniel

## Abstract

Decision-making in noisy, changing, and partially observable environments entails a basic tradeoff between immediate reward and longer-term information gain, known as the exploration-exploitation dilemma. Computationally, an effective way to balance this tradeoff is by leveraging uncertainty to guide exploration. Yet, in humans, empirical findings are mixed, from suggesting uncertainty-seeking to indifference and avoidance. In a novel bandit task that better captures uncertainty-driven behavior, we find multiple roles for uncertainty in human choices. First, stable and psychologically meaningful individual differences in uncertainty preferences actually range from seeking to avoidance, which can manifest as null group-level effects. Second, uncertainty modulates the use of basic decision heuristics that imperfectly exploit immediate rewards: a repetition bias and win-stay-lose-shift heuristic. These heuristics interact with uncertainty, favoring heuristic choices under higher uncertainty. These results, highlighting the rich and varied structure of reward-based choice, are a step to understanding its functional basis and dysfunction in psychopathology.

## Introduction

Agents making sequential choices under uncertainty face a pervasive tradeoff between learning about their environment and capitalizing on known rewards. This “exploration-exploitation” dilemma applies to bumblebees deciding where to forage ^1^, children making causal inferences ^2^, adults deliberating economic choices ^3^, and algorithms running stochastic optimization ^4^. Under most ecological conditions, there is either no known or no practical optimal solution to this tradeoff, in the sense that it would provide theoretical guarantees that long-term rewards are maximized ^5^. Such ecological conditions arise when observations are noisy, reward contingencies change over time, the decision problem has a long or indefinite time horizon, and/or the decision space is large, among others. A broad question that arises, then, is what are the algorithms that organisms, and humans in particular, use to resolve the explore-exploit tradeoff in practice and make sensible decisions?

Based on Gittins’ ^6^ original finding that an optimally exploring agent (in a particular simple and mathematically well-characterized setting) ought to confer an “uncertainty bonus” to options in addition to their expected reward values ^5^, a number of modeling studies have explored the role of uncertainty in helping humans balance the tradeoff between exploration and exploitation. Some studies have found support for these or similar strategies in humans ^7–12^, but there remain outstanding puzzles. First, results are rather mixed, with some other studies reporting uncertainty-avoidance effects ^13,14^, or no uncertainty-based exploration effects ^15,16^. This raises the question of what may cause heterogeneity in the use of uncertainty-based exploration strategies. Previous work has found substantial individual differences in the use of uncertainty in exploration, linked to psychological measures including impulsivity and anxiety ^15,17–20^. Here, we tested the hypothesis that there is considerable heterogeneity of uncertainty effects that reflects differences across individuals. Such heterogeneity could explain mixed results of previous studies, especially in small or non-random samples.

Second, some apparent effects of uncertainty can be accounted for by more basic strategies than incorporating uncertainty in decisions trial-by-trial. In many experimental contexts, such as ones involving stationary rewards and short time horizons ^7–9,11,17^, simple proxies for uncertainty, such as familiarity,novelty, or count-based heuristics may drive exploratory choices ^7,13,21,22^. Other basic heuristics, which do not require integrating information over many trials, such as a repetition bias or a win-stay-lose-shift strategy, may also mimic uncertainty effects, if not controlled for ^23,24^. For example, a tendency to switch upon encountering a large negative deviation from expectation to an expectedly lower value option can appear mistakenly as uncertainty-seeking behavior. Likewise, repetition bias, a form of perseveration (the tendency to repeat an option even when this does not maximize immediate reward), can appear uncertainty-driven, depending on how the decision policy is parameterized. Such basic heuristics are ubiquitous in human and animal decision-making ^25–35^, and may reflect cognitive constraints on decision-making ^36–40^. This raises the question of whether uncertainty effects are still observed when these heuristics are taken into account. It also raises the possibility that uncertainty may play further roles in human decision-making by interacting with such basic heuristics, i.e., by increasing or decreasing the degree to which they are used in a context-specific manner. In the present study, we set out to test potential roles of multiple decision factors, by identifying the contributions of these basic heuristics, repetition and wins-stay-lose-shift, alongside uncertainty and expected reward in guiding sequential choices, and to investigate their potential interactions.

We used a novel task design that deconfounds uncertainty from alternatives and expected reward. To anticipate our results, we find that (1) multiple factors in addition to uncertainty contribute to subjects’ choices; (2) uncertainty has heterogeneous effects on choices across subjects, and (3) these differences are stable across testing days within an individual; (4) repetition bias and win-stay-lose-shift tendencies contribute to choices pervasively across subjects; and, finally, (5) uncertainty modulates the use of these heuristics.

## Results

### 3.1 A novel explore-exploit task to characterize the role(s) of uncertainty

To study the roles of uncertainty in choices within a broader set of factors implicated in reward-based decisions, we adopt a novel variant of a two-armed bandit task (Fig. 1A). Subjects make a long sequence of choices (96 per block) between two options delivering rewards between 1 and 100 points with independently evolving reward distributions. They are only shown the reward of the chosen option, creating an explore-exploit tension. The need to explore is maintained by abrupt uncued shifts in reward levels (one out of three possibilities), termed changepoints (which occur independently for the two options), and noisy observations on each trial (drawn from Gaussians around a mean reward level). Cued low and high levels of observation noise are used across halves of a block to make changepoints more or less detectable, respectively, and to add further variation in estimation uncertainty (i.e. the observer’s uncertainty about the latent reward level; thereafter simply termed “uncertainty”). Expected reward and uncertainty tend to be negatively correlated in bandit tasks with partial information, because more rewarding options are (rightly) sampled more often ^11^. To decorrelate them, we interleave forced choice periods with free choices (see Methods for more details). A notable advantage of using long, nonstationary sequences, is that neither novelty nor familiarity ^7,13,21^ can account for exploratory choices, providing a more compelling test for genuinely uncertainty-based components of choice. Finally, subjects are occasionally asked to make explicit reports of reward level estimates and uncertainty / confidence about these guesses, to provide a choice-independent and direct (model-agnostic) measure of the factors that may guide their choices.

**Figure 1.**
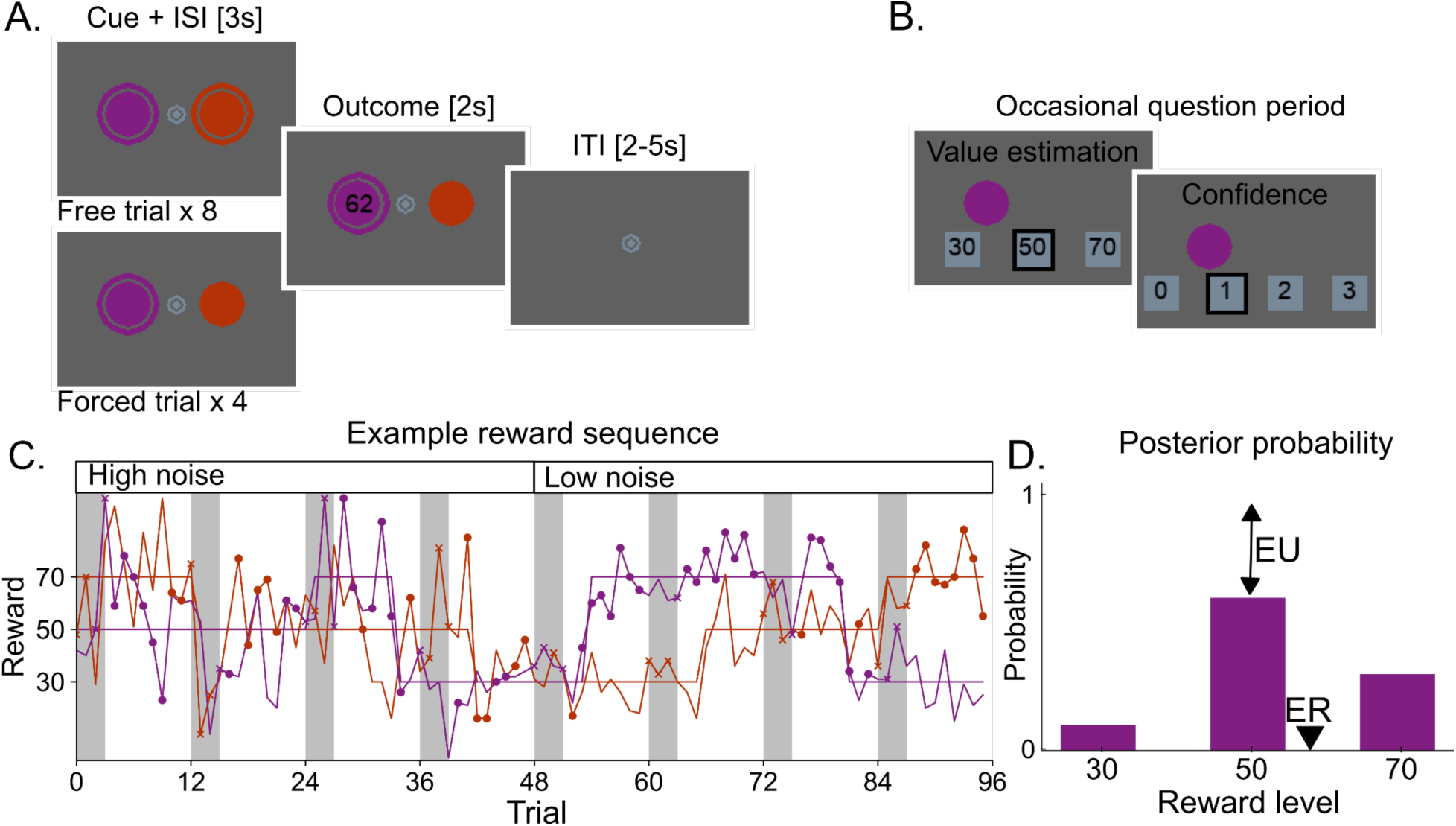
Task and model. **A.** On each trial subjects choose between two options and observe the outcome of the selected option. On forced trials, only the circled option can be selected (here, the purple one). **B.** Every 16 trials on average (SD=4.85) trials, subjects are asked to report their estimate of option values (black outline). Example of reward level estimate (left) and confidence report (right); only one option is shown here but both options are queried sequentially **C.** Example block. The latent mean reward levels (solid lines) shifted abruptly between 3 reward levels at uncued changepoints, independently for each option. Outcomes (dots) were drawn from Gaussians around the mean level with either low or high standard deviation. Each block was split into low-noise and high-noise halves corresponding to the low/high standard deviation of sampled outcomes. Subjects were instructed about noise levels, which were cued using a single or double circle around the fixation dot. Periods of 8 free trials alternated with periods of 4 forced trials (denoted by white and gray rectangles, circles and crosses, respectively). **D.** Posterior probability of an option following a single observation computed by the Bayesian ideal observer model. Model-based quantities were derived from the posterior: Expected reward (ER) is the sum of the reward levels weighted by their probabilities. Estimation uncertainty (EU) is 1 minus the probability of the maximum a posteriori (MAP) reward level.

We use a Bayesian ideal observer as a learning model to derive candidate decision factors. This learning model estimates the posterior probability of the latent reward levels for each option given the choices and outcomes observed by subjects. Notably, unlike point-estimates of reward expectation obtained from standard delta-rule models, the Bayesian learner also provides trial-by-trial uncertainty estimates about the latent reward level, which can be used as a proxy for subjects’ uncertainty ^41^. This formalization of uncertainty corresponds to *estimation uncertainty*, which depends both on the noisiness of the observations (variously known as risk, outcome variability, irreducible or expected uncertainty; ^14,42,43^) and the sampling history. The decision factors that we derive from the learning model are (in addition to a *repetition bias*): *immediate reward*, defined as the relative expected reward (ΔER; difference between the two options); the *relative and total uncertainties* (ΔEU, EUt; corresponding respectively to the difference and sum across the two options), both of which have previously been implicated in uncertainty-based exploration ^8,9,15^; a pair of factors capturing a *win-stay-lose-shift heuristic* (PE, UPE; the effect of signed and unsigned prediction errors on choices capture respectively the win-stay-lose-shift behavior and its potential asymmetry between the “win” and “loss” domains); and interactions between the uncertainties and the remaining factors (see Methods).

Our analysis strategy is as follows. Two prerequisites are, first, to validate that the Bayesian learning model accounts for subjects’ choices and reports, and second, to select the most appropriate form of choice stochasticity in the decision model (softmax or purely random decision noise). The learning and decision (i.e., policy) components are considered sequentially here for exposition, but in practice they are necessarily modeled jointly. The learning model can only be evaluated under a particular policy, and conversely, the definitions of the variables that comprise the decision policy depend on the learning model. Throughout this paper, we model the choice to repeat the previous decision or switch (rather than choosing option A vs B), because it provides a natural way to capture repetition bias, which we find to be a major contributor to choices in the task (see Results).

With these prerequisite modeling choices settled, our first main goal is to determine which factors contribute to choices via a model selection procedure. We then characterize the contributions of the main drivers of choice in addition to immediate reward, focusing first on the uncertainty-based factors and then on the basic choice heuristics, repetition bias and win-stay-lose-shift. Again, these sets of factors are considered sequentially for exposition, but they are modeled jointly, to determine each factor’s contribution while controlling for the rest. To test the hypothesis that variability in uncertainty effects across subjects reflects stable and psychologically meaningful individual differences, we assess the stability of uncertainty coefficients across testing sessions and their correlations with psychometric scales. To test the hypothesis that uncertainty modulates the use of the other heuristics, we ask whether and how uncertainty interacts with the remaining decision factors.

### 3.2 Task performance

Firstly, we verified the subjects understood and were engaged in the task. Out of 59 subjects, only 3 (5%) did not perform significantly above chance (49.5 points, SD=0.56). These 3 subjects were excluded from further analysis (the main conclusions remain unchanged when they are included). For the remaining 56 subjects in the final sample, the average reward earned was 54.0, SD = 1.16; range 51.2 - 55.6 points. This performance was on average slightly but reliably lower than simulated optimal performance under the Bayesian learning model and the selected decision model presented below (see Section 3.5) (M = 55.1; SD = 0.05, p=10^−8^). The overall commensurate performance in subjects and in simulation suggests that the model adequately captures subjects’ performance in the task.

### 3.3 A Bayesian model of learning accounts for choices and reports

#### Choices

It is necessary to assume a learning model to study the latent factors that guide choices and their values on a trial-by-trial basis. A Bayesian model has an *a priori* advantage for studying the roles of uncertainty in decisions: it provides trial-by-trial uncertainty estimates, which can be used as a proxy of subjects’ uncertainty.

However, simpler reinforcement learning models, such as the Rescorla-Wagner (RW) delta rule model, are commonly used ^44–47^. In contrast to a Bayesian model, this standard RW model only provides point estimates (no estimate of uncertainty), and assumes subjects do not use a learning procedure that is sensitive to the generative structure of the task. The standard version of RW only updates the expected reward of the option, for which there is a prediction error (i.e. the selected option here). To improve the ability of RW to better account for subjects’ choices, we augment it with updates of the unobserved option toward the average reward level, a feature that is also present in the Bayesian model (see Methods). The augmented RW model provided a better fit to choices than the standard RW (cross-validated average choice likelihoods for augmented RW: p_cv_(choice) = 0.702; for standard RW: p_cv_(choice) = 0.694; paired difference: 0.007, SE=0.0007, *t*(55) = 9.95, *p* = 10^−15^, Cohen’s *d* = 1.34). We find the Bayesian learning model accounts for subjects’ choices better than this augmented RW model (for the Bayesian model: p_cv_(choice) = 0.708; for RW: p_cv_(choice) = 0.702; paired difference: 0.007, SE=0.002, *t*(55) = 4.57, *p* = 10^−6^, Cohen’s *d* = 0.64; see Methods).

#### Reports

The inclusion of occasional explicit reports of the inferred mean reward levels and confidence permits an independent test of whether subjects represent value and uncertainty, and whether these representations resemble those of the Bayesian learner. Subjects’ guesses of the latent reward level match the true generative reward level and even more the ideal observer’s maximum a posteriori estimate well above chance (Fig. 2B) (fraction of reports matching generative values: 0.58, SE=0.012; matching the ideal observer: 0.61, SE=0.015; comparison to chance level (0.33): *t*(55)>19.20, *p<*10^−27^, Cohen’s *d* = 5.63; paired difference: 0.033, *t*(55) = 4.16; *p* =0.0001, Cohen’s *d* = 0.57), suggesting that the Bayesian latent reward estimates capture subjects’ subjective reward estimates.

**Figure 2.**
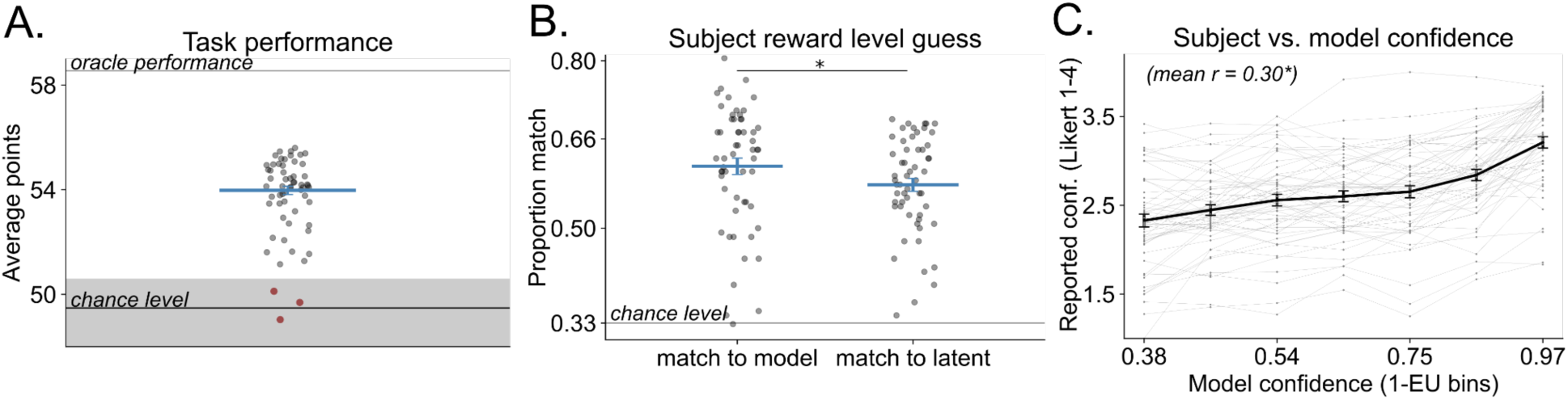
Choice performance and reports. **A**. Average performance (i.e. payoff) across sessions. Oracle performance is the expected payoff when selecting the option with highest (unknown) latent value on each trial. Chance performance is the expected payoff when selecting randomly and evenly both options. **B**. Fraction of reports matching the Bayesian model estimate of the reward level or matching of the (unknown) latent reward level. **C**. Average subject confidence for bins of Bayesian model confidence (with volatility fitted to subjects’ choices) about the latent reward level (1-EU). In all panels: dots correspond to subjects, error bars to SEM.

We also find subjects’ reported confidence reliably correlates with the Bayesian learner confidence (quantified as 1-EU) on the same trial (Fig. 2C): Pearson’s r_mean_ = 0.30, SE = 0.02, *t*(55) = 14.60, *p* = 10^−19^, Cohen’s *d* = 1.97. At the individual level, this correlation is significant at *p* < 0.05 in 77% of subjects (43 of 56). This suggests that subjects track uncertainty, and do so in a way that is similar to the Bayesian model. These results converge with evidence from behavioral modeling that the Bayesian model is a suitable model of subjects’ learning in the task.

#### Fitting volatility

Learning in the Bayesian model depends on the assumed volatility (probability of a changepoint on each trial). Analysis of this volatility parameter, fitted to each subject, provides convergent support that the Bayesian model is a useful approximation of subjects’ learning. Prior work suggests that humans tend to overestimate volatility ^20,48–50^. Consistent with this, the volatility parameter fitted in each subject was on average higher than the generative volatility (vol_fitted_= 0.16, SE=0.01, *t*(55) = 11.33, *p* = 10^−15^; vol_generative_ = 0.042, one-sample t-test against vol_generative_; Cohen’s *d* = 1.12), and models with volatility as a subject-specific free parameter outperformed equivalent models using the generative volatility (average difference in choice likelihood across models Δp_cv_ = 0.02, SE = 0.002, *t*(55) = 9.83, *p* = 10^−12^, Cohen’s *d* = 1.31). Therefore, all reported results are from models with fitted volatility.

Further validating the Bayesian learning model, we compare the correlation of subjects’ explicit confidence reports to the model confidence with (Fig. 2C) vs without (not shown) fitting volatility. The confidence estimates derived from the model with fitted volatility were more strongly correlated with subjects’ ratings than those derived from the generative volatility model: Δr = 0.03, SE=0.009, r_fitted_vol_ = 0.30, r_generative_vol_ = 0.26, *t*(55) = 3.89, *p* = 10^−4^, Cohen’s *d* = 0.53. The Bayesian estimates of the posterior reward level with the generative vs. fitted volatility did not differ in how well they match the explicit reports (p(match)_fitted_vol_ = 0.613; p(match)_generative_vol_ = 0.618, *n.s.*). This may be due to lower sensitivity of this measure, relative to correlations with confidence ratings. These results suggest that by estimating the subjects “presumed volatility” from choices, the Bayesian learner provides an even better approximation to the learning process, which is reflected in a closer match of the model to subjects’ subjective confidence.

### 3.4 Choice stochasticity is better captured by a reward-guided than a fully random policy

Models of choices typically assume some degree of choice randomness to handle residual, unexplained variance in choices. It is relevant to distinguish between two forms of randomness in the context of exploration. Epsilon-greedy decision models postulate that choices either strictly maximize the decision value or are made randomly with a probability epsilon on each trial. The random component of such a model captures completely random exploration. In contrast, softmax decision models postulate that choices maximize the decision value more often when the estimated reward value difference is larger between options. This random component corresponds to another form of exploration that is not completely random, but inversely related to the value difference.

To determine which form of choice randomness better accounts for our data, we compare cross-validated model fits with a softmax policy and an epsilon greedy policy, for the model including only a repetition bias and expected reward. The softmax policy provides a better fit to choices than epsilon greedy: base model (i.e., the model including all main effects, and no interactions): softmax p(choice)_cv_ = 0.708, ε-greedy p(choice)_cv_ = 0.650; Δp_cv_ = 0.058; SE = 0.007, *t*(55) = 7.99, *p* = 10^−10^, Cohen’s *d* = 1.08. Our interpretation of this result is that subjects’ propensity to choose the higher-valued option depends on the value difference. All reported results therefore use the softmax model.

### 3.5 Multiple factors contribute to choices, with a dominant effect of expected reward

Having validated the Bayesian learning model and selected softmax as the model of choice randomness, our first main goal is to determine which factors drive behavior in the task. Subjects were instructed that their goal was to maximize points earned in the task, and the observed task performance (Section 3.1) already implies that immediate reward is a strong driver of choices since subjects are, on average, more likely to choose the more rewarding option. Our model selection strategy is therefore to consider a baseline model, consisting of the immediate reward term and a repetition bias (as the intercept), and to ask what additional factors significantly improve the cross-validated model fit, including the relative and total uncertainty (ΔEU and EUt, respectively capturing uncertainty-based exploration), and two prediction error terms (signed and unsigned PE, capturing the win-stay-lose-shift heuristic). We iteratively add all combinations of one to four factors, for a total of 16 possible combinations. Model pairs which only differ in the inclusion or exclusion of a single factor showed that each of the four additional factors significantly contributes to choices (Fig. 2A). To ensure that the parameters of the full model are distinguishable and that choices generated with a given model are best fit by the same model, we performed parameter and model recovery (Fig. S1 and S2).

A strong test for the individual contributions of each factor are single-factor ablations from the full model, testing for the marginal contribution of a given factor while accounting for the rest (Fig. 2A, bottom row). The respective unique additional contributions of each factor in this test are Δp_cv_(choice): ΔEU = 0.11% (SE = 0.03, *t*(55) = 4.14, *p* = 10^−4^, Cohen’s *d* = 0.56); EUt = 0.07%, SE = 0.03, *t*(55) = 2.67, *p* = 10^−3^, Cohen’s *d* = 0.36); PE = 0.47%, SE=0.11, *t*(55) = 4.15, *p* = 10^−4^, Cohen’s *d* = 0.56, UPE = 0.19% (SE=0.04, *t*(55) = 4.65, *p* = 10^−4^, Cohen’s *d* = 0.63). Together, these four factors contribute an extra percentage-point (1%) to the cross-validated choice probability (p_cv_(choice) in the full model: 0.718, in the base model: 0.708). This should be compared to the respective unique additional contribution of expected reward (7.28%, SE = 0.45, *t*(55) = 16.26, *p* = 10^−21^, Cohen’s *d* = 2.17) and of the repetition bias (5.72%, SE = 0.71, *t*(55) = 8.05, *p* = 10^−10^, Cohen’s *d* = 1.08) in the full model. The above-chance performance of the model without the dominant factors (ablate ΔER: p(choice)_cv_ = 0.65; ablate repetition bias: p(choice)_cv_ = 0.66; ablate both: p(choice)_cv_ = 0.59) reflects shared predictive variance among decision factors.

Model comparison within subjects indicates that the full model is the most frequent best model at the subject-level, followed by the two models that differ from the full model by lacking only either ΔEU or EUt (Fig. S3). This is an important result that rules out that the full model is the best at the group level because it is the only one that can accommodate the implication of different decision factors among subjects. Instead, it is the best model because many factors (including uncertainty) contribute to choices within subjects.

We also note that ignoring the heuristic factors (repetition bias, or PE and UPE) from the model of subjects’ choices inflates the estimate of ΔEU (Fig. S4); accounts that omit the contribution of these heuristics may thus lead to erroneous conclusions regarding uncertainty.

### 3.6 Uncertainty has subject-specific effects on choices

#### Heterogeneity in uncertainty effects across individuals

The model accounts for the potential effect of two aspects of uncertainty on choices. ΔEU captures either uncertainty-seeking (analogous to an uncertainty bonus ^7,9,51^) or uncertainty avoidance, depending on the sign of the coefficient (w_ΔEU_). EUt captures a “default” choice strategy to repeat the previous choice or switch when overall uncertainty is higher (depending on the sign of the coefficient, w_EUt_). The coefficient estimates are on average close to zero, with a slight uncertainty avoidance effect (w_ΔEU_ = −0.073, SE = 0.028, *t*(55) = −2.57, *p* = 0.01, Cohen’s *d* = 0.34) and a slight tendency to repeat the previous choice when overall uncertainty is higher (w_EUt_ = 0.053, SE = 0.023, *t*(55) = 2.33, *p* = 0.02; Cohen’s *d* = 0.31. Fig. 2B; these parameter estimates are from the full model with interactions, see Fig. S5 for the model without interactions). These results suggest that the heterogeneous contributions of these variables across individuals, which improve the model fit, manifest as null or small regression coefficients on average at the group level.

#### Uncertainty effects capture stable, psychologically meaningful individual differences

To test the hypothesis that the inter-subject variability of uncertainty-related parameters reflects true differences between individuals rather than noise in the data, we assess the stability of the coefficient estimates within subjects across time. We fit separate models to each subject’s data from the behavioral sessions on the one hand and fMRI sessions on the other hand, and compute the Spearman correlation of the resulting parameter estimates. The sessions were completed on different days, and with differences to adapt the task for the MRI scanner (see Methods). As a useful point of comparison, the coefficients of the strongest decision factors (repetition bias and ΔER) are highly correlated across sessions (repetition bias: *r* = 0.658, p = 10^−7^; ΔER: *r* = 0.618, *p* = 10^−6^; Fig. 3A, bottom), as are the coefficients for the win-stay-lose-shift heuristic terms: PE (r = 0.463, *p* = 10^−4^), UPE (r = 0.450, *p* = 10^−3^). Critically, the ΔEU (*r* = 0.49, *p* = 10^−4^) and, to a lesser extent, the EUt (*r* = 0.20, p = 0.13) effects are also stable across sessions. A qualitatively similar pattern is obtained for the full model with interactions, although in the fMRI session, the interaction terms are less reliably estimated than in the behavioral session, probably reflecting the two-fold difference in number of trials between sessions (Fig. S6). These results suggest that heterogeneity in the uncertainty effects reflects persistent individual differences rather than noise.

**Figure 3.**
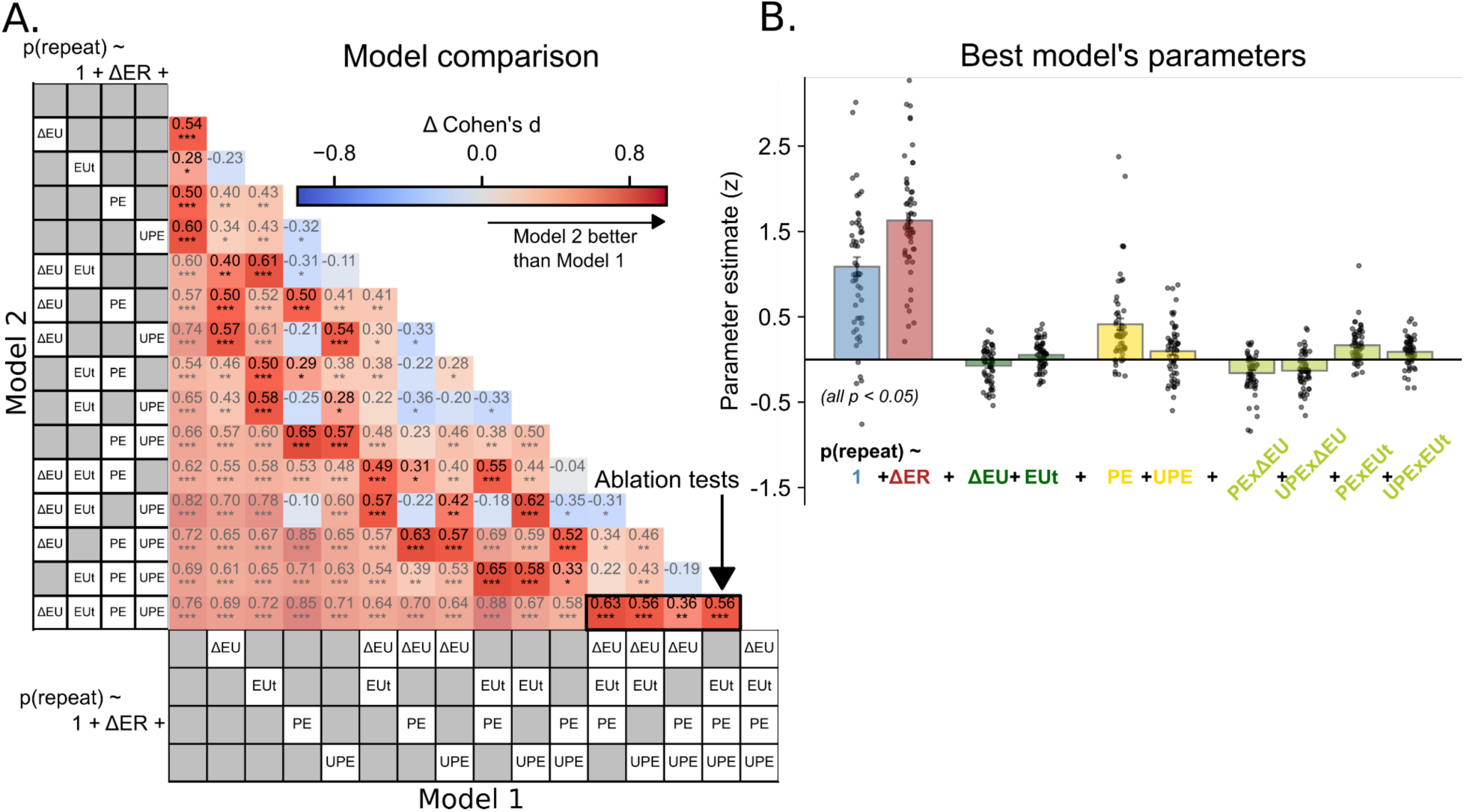
Multiple factors contribute to choices. **A**: Model selection of main effects. All considered decision factors contribute to choices. Adding 1-4 factors consecutively to a base model of repetition bias + ΔER improves the cross-validated model fit (average choice likelihood). Colors represent effect sizes (Cohen’s *d*) of model fit difference. Highlighted squares (in contrast to semi-transparent ones) are minimal model pairs with a one-factor difference, showing reliable improvements from adding each term. The bottom row and four rightmost columns (Ablation tests) show unique factor contributions controlling for all other factors; *: p<0.05, **: p<0.01, ***: p<0.001 **B.** The selected (full) model coefficient estimates (all *p* < 0.05; dots are individual subjects) show expected reward (ΔER; red) and repetition bias (blue) to be the dominant drivers of choices, with heterogeneous uncertainty effects (ΔEU and EUt; green) and an additional contribution of the signed and unsigned previous prediction error (PE and UPE, yellow), capturing a win-stay-lose-shift (WSLS) heuristic. Uncertainty interacts with the WSLS heuristic (light green bars), such that when relative uncertainty is higher on the previously unchosen option or total uncertainty is higher, subjects relied more on the heuristic.

Next, we test whether these stable individual differences track with psychometric scales that measure psychological dimensions. Corroborating previous work showing that anxiety is associated with uncertainty avoidance in a bandit task ^19^, individuals who score higher on anxiety are more likely to avoid the more uncertain option (ΔEU, estimated on all data; state anxiety: Pearson’s *r* = −0.32, *p* = 0.016; Fig. 3B; trait anxiety: *r* = −0.23, *p* = 0.08). A similar but weaker result is obtained for ΔEU coefficients using the full model with interactions (Fig. S7). Previous work has also found that compulsive gambling, associated with impulsivity, is related to stronger information-seeking in a bandit task ^17^. In the present data, more impulsive individuals also tend to show greater uncertainty-seeking (Pearson’s *r* = 0.22, *p* = 0.09; Fig. 3B), in the full model including interactions (see Fig. S7 for full correlation matrix of parameter estimates with psychometric scales).

Taken together, these results indicate that uncertainty plays a role in choices, with subject-specific, psychologically meaningful effects across individuals, which relate to trait measures related to anxiety and impulsivity.

### 3.7 Other decision heuristics drive choices

The individual differences in uncertainty effects we identify above can explain why previous studies have not always found a consistent uncertainty-seeking or uncertainty-avoiding tendency. In addition, some decision heuristics are partly confounded with uncertainty, and often not controlled for, which can also appear as uncertainty effects across studies. Specifically, in tasks with noisy observations, repeating the same choice typically reduces the uncertainty about the corresponding option, which could motivate the repetition. However, repetition may also simply reflect perseveration. This is a general behavioral tendency observed across many task contexts, which leads, in the context of reward-based choice, to selecting an option even when it is no longer the most rewarding one. Similarly, positively surprising observations – ones that generate large positive prediction errors – may result in repeating an option even when there is insufficient evidence that this option is most rewarding. Conversely, and perhaps more commonly, large negative prediction errors may cause switching to a less well-known option even when its expected reward is lower than the current option. When a win-stay-lose-shift strategy is not controlled for, these patterns may appear as uncertainty-seeking.

Consistently with these possibilities, we find that both repetition bias and terms that capture a win-stay-lose-shift strategy (the signed and unsigned previous prediction error, PE and UPE) significantly improve the model fit, after controlling for immediate reward and uncertainty-based effects (see Section 3.5). Furthermore, the coefficients for these effects show a consistent direction at the group level (while accounting for other factors): a bias to repeat rather than switch: w_0_= 1.087, SE= 0.111, *t*(55) = 9.76, *p* = 10^−12^, Cohen’s *d* = 1.30 and win-stay-lose-shift decisions: w_PE_ = 0.412, SE = 0.069, *t*(55) = 6.02, *p* = 10^−6^, Cohen’s *d* = 0.80. A follow-up analysis of UPE shows that this term accounts for an asymmetry in win-stay-lose-shift decisions: subjects are more likely to repeat given large positive prediction errors than to switch in response to similarly large negative prediction errors (Fig. S8). Interestingly, the heuristic effects are also individually stable across sessions (Fig. 4A, bottom) and exhibit meaningful correlations with psychometric scales (e.g. between w_PE_ and optimism; between w_0_ and anxiety and impulsivity, see Fig. S7).

**Figure 4.**
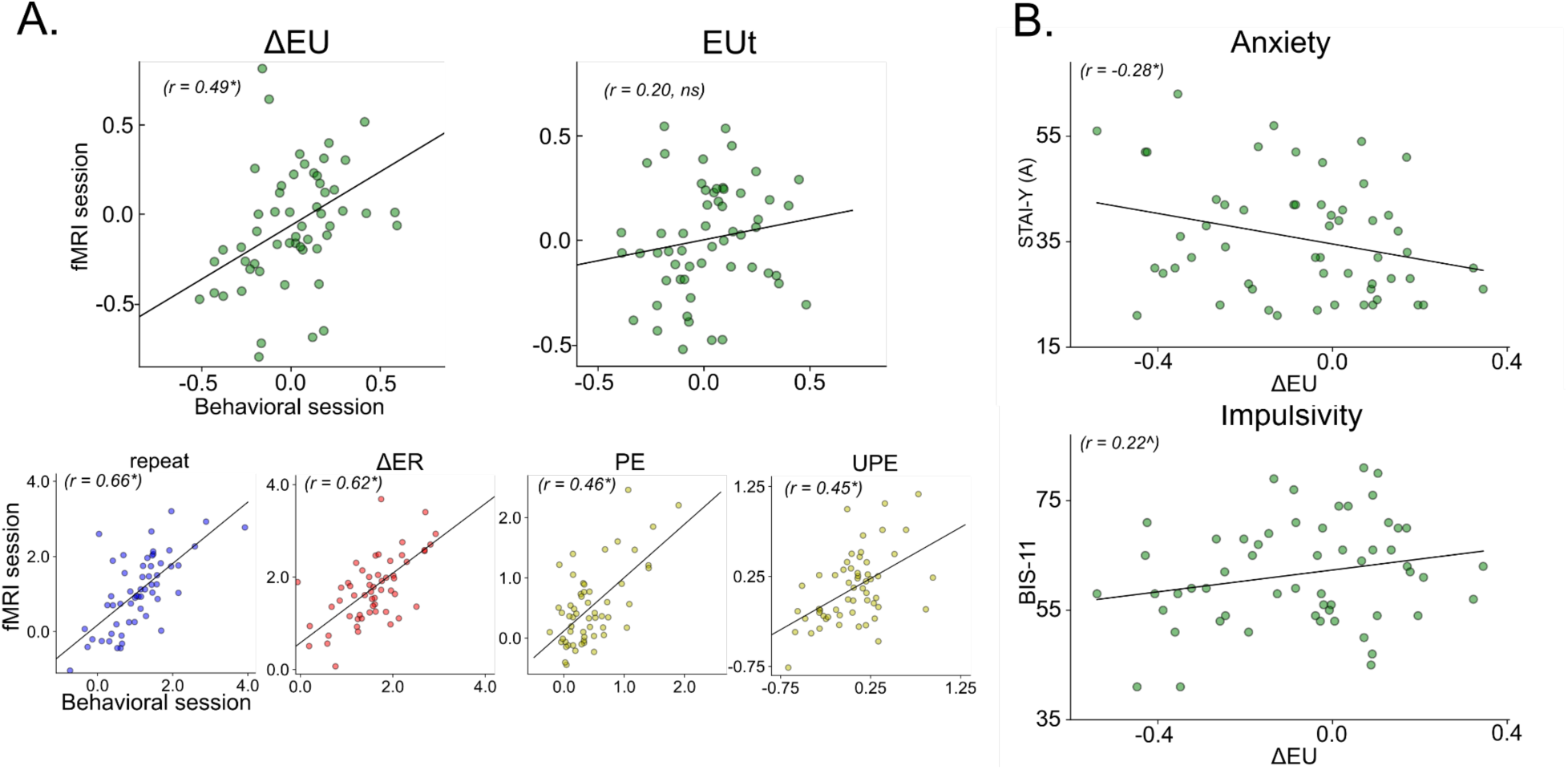
Effects of uncertainty on choices are subject-specific. **A. Top**. Scatterplots of regression coefficients estimated per session (behavioral and fMRI) show stability of uncertainty estimates. Dots are subjects. **Bottom**. Other model factors are also correlated across sessions. **B.** Uncertainty effects (ΔEU) correlate with psychological measures of anxiety and impulsivity. *p < 0.05; ^0.05 < p < 0.1.

In sum, we find that basic decision heuristics, which are often not controlled for in studies of exploration, consistently contribute to choices across subjects.

### 3.8 Uncertainty modulates the use of the other heuristics

Heuristics are decision rules that subjects may be more likely to use when evidence about value is scarce or similar across options. In other words, uncertainty may modulate the extent to which behavior depends on heuristic decisions. To test this hypothesis, we first perform a two-step model selection analogous to that for the main effects model, but focused on testing interactions with uncertainty.

First, starting from a base model including all main effects, we add two-way interactions of each of the two uncertainty terms, ΔEU and EUt with either ΔER, or PE and UPE (together because they jointly capture the win-stay-lose-shift heuristic) (Fig. 3B and Fig. S8).

The result of this first step is that neither of the interactions with ΔER (ΔER*ΔEU and ΔER*EUt) individually contribute significantly to the model fit, they are therefore excluded (p_cv_(choice) with ΔER*ΔEU: *t*(55) = −0.20, *n.s.*; p_cv_(choice) with ΔER*EUt: *t*(55) = 0.65, *n.s.*). Both of the interactions of the uncertainties with the win-stay-lose-shift heuristic individually improve the model fit, so they are included in the second step. In the second step, we test whether adding both of these (pairs of) interactions further improves the model fit over either one in isolation. It does, so both are kept in the final model. Again, the fit improvement in terms of additional variance explained over the main-effects model is modest but reliable, with moderate effect size (0.17%, SE = 0.04, *t*(55) = 4.41, *p* = 10^−4^, Cohen’s *d* = 0.59); the fit improvement over the base model (intercept and ΔER) is comparable to that of adding the additional main effects (1.10%, SE = 0.18, *t*(55) = 6.29, *p* = 10^−7^, Cohen’s *d* = 0.84). There is therefore sufficient evidence that these interactions should be included in the model (Fig. S9).

To characterize their contributions, we examine the coefficient estimates for the interactions (Fig. 3B). When EUt is higher, subjects tend to rely more on the win-stay-lose shift heuristic. When the relative uncertainty, ΔEU, of the previously unchosen (switch) option is higher, they also rely more on this heuristic. Recall that the main effect of EUt, presented above, captures a tendency to repeat more when overall uncertainty is higher, which can also be seen as reinforcing a basic heuristic repetition bias. In sum, when faced either with more uncertainty across both options or more uncertainty in the unchosen option, subjects are more likely to base their choice on heuristics.

## Discussion

In this study, we used a novel bandit task to simultaneously characterize the contributions of uncertainty and other factors in reward-based decisions. We found a dual role for uncertainty in choices. First, uncertainty had subject-specific effects on choices, which were stable over time (correlated across testing sessions) and psychologically meaningful (correlated with psychometric measures of anxiety and impulsivity). Second, uncertainty modulated the use of two basic heuristics, repetition bias and a win-stay-lose-shift strategy: subjects rely more on these heuristics when overall uncertainty is higher and also when the other option (switch) is more uncertain than the current option (stay). Finally, the heuristics were themselves consistent drivers of choices: controlling for expected reward and uncertainty, subjects showed a general tendency to repeat rather than switch, and a tendency to overweigh the most recent observation, to repeat the previous choice in response to positive prediction errors and to switch in response to negative ones. However, it is important to note that, since the uncertainty-related coefficients were estimated in the presence of these heuristic terms, the subject-specific uncertainty-seeking/avoiding behavior is genuine.

These results provide a richer characterization of the interplay between value, uncertainty, and basic heuristic factors, where the latter are often neglected in studies of exploratory decision-making. They also inform an important puzzle in computational research on the exploration-exploitation dilemma: if uncertainty-based exploration strategies are generally an effective solution to explore-exploit tradeoffs, why are uncertainty effects not more robust and consistent across studies? The stable subject-specific uncertainty effects we found, ranging from uncertainty seeking to indifference and avoidance, point toward one part of an answer: that mixed results, especially from studies with smaller and non-random samples of subjects, may arise from underlying population-level heterogeneity in the uses of uncertainty. The heuristic decision factors we identify suggest another part of an answer. First, when not explicitly modeled, heuristic choices may appear either as exploratory effects or as uncertainty avoidance, depending both on the specific operational definition of exploration and on specifics of experimental design. Second, how individuals manage the balance between a plurality of decision factors in addition to reward and uncertainty may be a further source of heterogeneity. This balance is plausibly itself negotiated by recourse to uncertainty, as suggested by the uncertainty-by-heuristic interactions we report and by prior work ^52,53^.

These interpretations, elaborated below, recast the question of variable uncertainty effects at the individual level: if uncertainty-based exploration is essential to effective reward-based decisions, why do individuals vary in whether and how they use uncertainty? There are two compatible possibilities. First, the correlations of uncertainty with psychometric measures of impulsivity and anxiety suggest individual differences in how well people leverage uncertainty in making decisions, which in extreme cases may be related to psychopathologies. Second, there may actually be a normative, resource-rational basis ^39,40^ for the individual differences observed in our study and elsewhere.

A growing number of studies have related individual differences in responses to uncertainty and exploratory behaviors to psychometric scores or clinical diagnoses, particularly of anxiety and impulsivity, consistent with our findings. With regard to impulsivity, a recent large-sample study of the general population has shown increased random exploration in more impulsive individuals, but no specific relationship to uncertainty-based exploration ^18^. Two studies of patients with gambling disorder ^17^ and attention deficit hyperactivity disorder ^54^, which are thought to involve impairments in impulse control, have respectively found increased information seeking and random exploration. Although these studies do not speak to the direct role of uncertainty in choices, they are broadly consistent with the present results in suggesting increased exploratory tendencies in more impulsive individuals. Interestingly, another study of gambling disorder ^55^, found decreased uncertainty-directed exploration in gamblers, contrary to our results. This study used continuously drifting latent values^16^, rather than changepoints; this difference may be significant given that impulsivity might be specifically related to increased novelty-seeking ^56,57^ (cf. ref ^17^). Over the long trial sequences in both tasks, none of the options is truly novel, but abrupt shifts in reward levels in our task might effectively induce a novelty-like effect.

With regard to anxiety, perhaps the strongest evidence for a negative relationship between anxiety scores and uncertainty-directed choices in line with our results comes from a recent study by Fan *et al* with large online samples, which found reduced uncertainty-directed exploration (as well as uncertainty avoidance) in a bandit task^19^ (Fan et al., 2022). Notably, estimation uncertainty in the study by Fan *et al*, unlike ours, includes a component of outcome uncertainty differences across options, i.e., risk, leaving open the question of whether anxiety is related to risk aversion, as also suggested by another study ^58^, or estimation uncertainty. Our study adds to these results that more anxious individuals tend to avoid uncertain (i.e. lesser known) options even when risk levels are matched.

Another study found that in a non-instrumental context, when uncertainty reduction does not bear on choosing a more rewarding option, more anxious individuals were willing to pay a premium to resolve uncertainty ^59^, which in a bandit setting might suggest uncertainty seeking rather than avoidance. A general aversion to uncertainty characteristic of anxiety ^60^, is consistent with either action to reduce uncertainty or with uncertainty avoidance, and the difference in whether uncertainty is instrumental or intrinsically valued is plausibly significant ^21^. One recent study has, however, reported increased instrumental uncertainty-directed exploration with anxiety, contrary to our results ^61^. The interpretation of this other study is complicated by the use of a proxy for estimation uncertainty that is potentially confounded with perseveration and choice stochasticity. In sum, although results are somewhat mixed, there are converging indications that exploratory choices are affected by anxious and impulsive traits, and a next step is to determine the precise nature of these effects, their relationship to uncertainty, and their causal origins.

We now turn to the other major conclusion of our study: choices are underpinned by multiple decision factors, including heuristic factors besides expected reward and uncertainty. The basic heuristic factors we identified, repetition bias and win-stay-lose-shift strategy, are naturally interpreted in terms of decision-making mechanisms that are commonly contrasted with reward-based choice, namely habitual and Pavlovian responses, respectively. Here, we provide an interpretation of these heuristics in these terms, while acknowledging the possibility of alternative interpretations.

The repetition bias may reflect a form of perseverative or habitual response, whereby the previous selection of an action increases the probability of selecting that action again. The probability of repeating may increase either because of a caching of previously rewarded stimulus-action associations ^52,62,63^, or independently from reward ^25–27,64^. In either case, the hallmark of habits is that they are fast and often efficient, but inflexible because of their decoupling from current rewards^65^. The caching mechanism is perhaps broadly analogous to the repetition bias in our study: once a subject has found the more rewarding option, they may choose it without referring to its expected value on every trial. However, computational studies of habit have recently emphasized that there is no necessary link between habits and values: actions may become habitual simply because they are repeated; it is incidental that the actions that tend to be repeated are usually rewarding ones ^36,64^. This second view entails a more thorough dichotomy between two independent choice generating processes: a habitual and a reward-based one.

The win-stay-lose-shift heuristic we identified can be interpreted as an automatic tendency to approach stimuli previously paired with reward and to avoid ones paired with punishment (where “punishment” in the present context would be construed in relative terms, as lower-than-expected reward). This may occur even when such approach/avoidance behavior conflicts with the instrumentally dictated action, such as when expected reward recommends staying with an option that yielded surprisingly low reward on the last trial. This automatic tendency can be interpreted as “Pavlovian” in that it is governed by stimulus-outcome associations that are independent from instrumental considerations ^29,66–68^. Innate Pavlovian tendencies to approach appetitive stimuli and avoid aversive ones can be hard to overcome, as shown in animal studies of associative learning ^29,68–70^. In humans, Pavlovian biases have been demonstrated in Go/NoGo paradigms to conflict with instrumental requirements to withhold action to obtain a reward or to act to avoid punishment ^32,66,67^. These studies argue for the independence of a Pavlovian choice generator, which, similar to habits, produces automatic responses to stimuli that are not coupled with action-outcome contingencies.

Thus, given the ubiquity of these basic heuristics in animal and human sequential decision-making, they can be plausibly construed as independent choice generators alongside more complex reward- and uncertainty-based strategies. Notably, however, alternative interpretations cannot be excluded. For example, it is possible that the apparent multiplicity in choice generative processes may be an artifact of model mis-specification: an inaccurate formalization of the value function or the learning process with respect to the true latent values guiding subjects’ choices may manifest as repetition bias or win-stay-lose-shift choices. Specifically, because the intercept term in the logistic regression (i.e., repetition bias) captures the mean tendency to repeat rather than switch, there is always the possibility that some residual repetitions are reward-driven from the point of view of a subject’s value estimation ^71,72^, but not according to the Bayesian ideal observer. Similarly, with regard to win-stay-lose-shift, higher-than-optimal discounting of past rewards, due to, say, attentional and working memory limits, may manifest as choices that overweigh the last observation ^23,73^.

Studies of the exploration-exploitation dilemma often focus on reward- and uncertainty-based drivers of choice without accounting for potential additional factors, such as perseverative or approach-avoidance responses. The present results call for consideration of the status of the basic heuristics with regard to exploration. On the broadest sense of exploration as any choices that do not maximize immediate reward ^16,74^, basic heuristic-driven choices that diverge from immediate reward maximization are exploratory. It is not clear to what extent similar heuristic factors are present across the gamut of bandit settings used in the experimental literature. Stationary or slowly changing latent rewards would seem liable to result in perseverative, repetition effects. Perceptions of low controllability resulting from high reward stochasticity or volatility seem likely to increase the incidence of Pavlovian win-stay-lose-shift responses ^66^.

Similarly, uncertainty-based definitions of exploration may confound a genuine effect of uncertainty with more basic heuristic factors, when these heuristics are not explicitly modeled. Such confounding may, on balance, go in either direction, mistakenly suggesting either uncertainty-seeking or uncertainty-avoidance, or either increases or decreases in choice stochasticity with uncertainty. There are many conceivable patterns of confounding, and they may help explain the heterogeneity of uncertainty-based exploration effects in the literature. Future work should systematically characterize the recruitment of different decision heuristics with variations in task parameters. When they are not directly of interest, they should at least be controlled for.

Lastly, the interactions we identify of uncertainty with the basic heuristics bear on the preceding discussion in several ways. Methodologically, they may further aggravate confounding when relevant decision factors are unaccounted for. Substantively, two notable implications of these interactions are that, first, they may point toward an uncertainty-based arbitration mechanism between strategies ^52,53^, and second, individual differences in the strength and direction of such interactions may be a further source of heterogeneity in choice patterns. A promising direction for evaluating these possibilities in future work is to move away from models of average factor contributions over entire choice sequences and toward modeling the dynamics of strategy selection with temporally resolved methods, both descriptively (for e.g., with HMM-GLMs ^75^) and algorithmically.

In conclusion, the present study has showcased the rich structure of human sequential reward-based decisions, and has identified stable and meaningful individual differences in the uses of uncertainty in choice, which can account for mixed results of prior work. These results underscore the importance of characterizing the interplay between reward, uncertainty, and a broader range of cognitively relevant factors for understanding the processes governing human choice under uncertainty, their functional basis and dysfunction in psychopathology.

## Methods

### 2.1. Subjects

Sixty-two adult volunteers with no self-reported history of psychiatric or neurological conditions participated in the study and received a fixed monetary compensation. Two subjects were excluded for only completing one of the two sessions of the experiment. One subject was excluded for an incidental finding of brain anomaly in the MRI. Three subjects were excluded for chance-level task performance. Chance level (mean and SD) was defined as the average obtained reward across 1000 iterations of random choices on the same reward sequences as subjects, and the exclusion threshold was defined as mean obtained rewards less than two standard deviations above chance: M_chance_reward_ = 49.5 points, SD_chance_reward_ = 0.56, cutoff =50.6; the 3 excluded subjects obtained 50.1, 49.0, and 49.7 points on average. The data from the remaining 56 subjects were analyzed (M_age_ = 25.9 years, SD_age_ = 6.5, range 18-44, 29 women, 27 men; 48 self-reported right-handed; all native French speakers). The study was approved by a national ethics review board (Comité de protection des personnes Ile de France III, approval #2018-A09195-50), and subjects gave informed consent prior to participating.

### 2.2. Main task and procedure

#### Sessions

Subjects completed two 2-hour long in-lab sessions: first, a behavioral session composed of 8 blocks of the task (as well as instructions and practice, see below), and an MRI session on a separate day, consisting of 4 blocks. The sessions were between 1 and 29 days apart (M_days_=10, SD_days_=6.5). Subjects were encouraged to take occasional breaks in-between blocks to maintain attentiveness. Between the two in-lab sessions, subjects completed an online battery of psychometric questionnaires, on the Gorilla Experiment Platform (https://gorilla.sc/), detailed below.

#### Task structure

We developed a novel version of a non-stationary two-armed bandit task, which allowed (a) dissociating reward-driven choices from other decision factors, and (b) better identifying uncertainty-based vs. heuristic-driven choices. A block of the task (Fig. 1A, C) consisted of 96 partial-feedback trials (i.e., subjects only observed the outcome for the chosen option, creating an explore-exploit tradeoff), with rewards ranging between 1 and 100 points drawn from Gaussians centered around 3 mean reward levels of 30, 50, and 70 points. There were uncued, independent changes of the reward levels for each option every 24 trials, on average (min: 9, max: 36), for a total of 4 changepoints per option per block (volatility, i.e. probability of a change point on a given option at any given trial, = 1/24; with a minimum of 9 trials in-between consecutive changes on the same option). There were interleaved segments of 4 forced-choice and 8 free-choice trials (each block starting with a forced choice segment), whose goal was to decorrelate expected reward and estimation uncertainty, which are otherwise anticorrelated (i.e., the more rewarding option is selected more often thereby reducing its estimation uncertainty; Fig. S10) ^11^. Further, each block consisted of a cued high-noise (SD=20 points) and a low-noise (SD=10 points) period of 48 consecutive trials (in counterbalanced order across runs). Notably, the level of noise for the two options was matched throughout, therefore the relative risk level or irreducible uncertainty was always matched between the two options. The goal of this noise-level manipulation was to induce different levels of estimation uncertainty across trials, on top of the differences induced by change points. Within a block, the average rewards and variability for each option were matched.

#### Reward sequences

A total of 8 fixed sequences were used to maintain comparability across subjects and sessions, for purposes of investigating individual differences in model parameter estimates across subjects as well as parameter stability within subjects across sessions. Even though the underlying rewards on the two options were “fixed”, the exact sequences of rewards observed by each subject and across sessions varied for the free-choice periods because of partial feedback. The forced choices were fixed for each sequence (i.e., all subjects were required to choose the same option on a given trial, and consequently they obtained the same reward), and were balanced for equal (each option sampled twice in a 4 trial segment) vs unequal (options sampled 1 and 3 times) information across options ^7^. In the behavioral session, subjects completed 8 blocks, with each block consisting of a single presentation of each sequence in randomized order. In the fMRI session, subjects completed a subset of 4 of the same 8 sequences.

#### Visual presentation

The two arms of the bandit, referred to as “options”, were presented as a filled purple and orange circle on a gray background, and outcomes for the chosen option were shown in black numerals inside the respective circle. The location of the purple and orange option (left vs right) was counterbalanced across blocks but was held constant within a block. Notably, in this design, a tendency to repeat the previous option irrespective of reward history conflates motor perseveration (making the same motor response) and choice perseveration (choosing the same option).

A second, larger concentric circle around the option signaled free- vs. forced-choice trials: when only one of the options was “outlined” with a second circle, subjects had to select that option; when both options were highlighted, subjects could freely choose an option. Upon option selection, the circle around the selected option disappeared to provide feedback that the response was registered, and then reappeared for 500 ms before the obtained reward was displayed. Fixation was indicated by a bullseye at the center of the screen (lighter-gray relative to the background), which also served to signal whether subjects were currently in a low- or high-noise period: 2 concentric circles indicated low noise; 3 circles indicated high noise. When the noise level switched halfway through a block, a yellow circle flashed around the fixation bullseye to alert subjects to the change.

There was no indication of reward history from past trials, the number of trials elapsed and remaining in a block.

Subjects were instructed to maintain central fixation, and the visual display of the options subtended roughly 2.5 degrees of visual angle (from the outer edges of the left and right options). Subjects wore noise-canceling headphones in both sessions (Behavioral session: Bose noise canceling headphones 700; https://www.bose.com/; MRI session: MR Confon, https://www.crsltd.com/tools-for-functional-imaging/audio-for-fmri/mr-confon/). In the behavioral session white noise was played over the headphones.

#### Trial timing

The timing differed across the behavioral and MRI sessions. Specifically, the behavioral session was quasi-self-paced, with the following fixed components: a 1 s interval between button press and outcome, a 2 s presentation of the outcome, and a 1 s inter-trial interval (ITI) consisting of a fixation-only display between the disappearance of the last trial’s outcome and the reappearance of the cues for the next trial.. By contrast, in the fMRI session, the entire trial duration was fixed as follows: 2 s cue display and response window, 1 s interval between cue and outcome display, 2 s outcome display, 2 to 5 s ITI (3.5 s on average). Therefore, the average stimulus onset asynchrony in the scanner was 8.5 s (range 7-10 s). If subjects failed to respond within the 2 s response window, a red fixation cross was displayed for the remaining duration of the trial (M=5.02 (1.07%), SD=6.36).

#### Responses

In the behavioral session, subjects indicated their choices on a standard QWERTY keyboard, with an index-finger button press of the left and right hand for the left and right options, respectively, using the “f” and “j” keys. In the scanner, subjects made their responses with an index- or middle-finger button press (left and right options, respectively), via a custom-made MR-compatible optic fiber button box with 5 buttons, developed in-house.

#### Explicit reports

Subjects were occasionally asked to report guesses for the reward level of each option (30, 50, or 70 points) and their confidence in these guesses (4-point Likert scale from 0 = “Not at all” to 3 = “Totally”, corresponding to “Pas du tout” and “Totalement” in French). A “Questions” screen signaled the beginning of an explicit report period. Subjects first answered both questions for one option and then for the other.. In the behavioral session, subjects indicated value guesses on a keyboard with their left hand (keys: “s”=30, “d”=50, “f”=70) and confidence with their right hand (“j”=0, “;”=3). In the fMRI session, they answered both questions with the right hand, with thumb, index, and middle fingers for value guesses and additionally ring finger for confidence (low to high reward level / confidence mapping from thumb through middle / ring finger). The order in which the options were queried was counterbalanced within a block.

In each behavioral block, there were a total of 6 reports (each one querying both options), 4 of which were on free trials and 2 on forced trials. In an fMRI block, there were 4 reports, 2 on free and 2 on forced trials. The total number of explicit reports across both sessions was 128.

The timing of the responses was self-paced in the behavioral session and with a 3 s response window in the fMRI session. In the fMRI session, the next question appeared once a response was registered, but the total duration of a report period was kept fixed by adjusting the duration of fixation display following the last report.

#### Instructions and practice

In the first in-lab session, prior to the main task, subjects received written task instructions in French that relied on a backstory to facilitate understanding of the task structure. Subjects were given an opportunity to ask clarification questions. The backstory involved commodity traders of apples and oranges in noisy and volatile market conditions (see Fig. S11 for the English translation of the instruction with the backstory). They then completed two practice blocks: first, a full block (96 trials) of an illustrated version of the task matching a backstory; and second, a short practice block of the task as presented with a simplified display during the main experiment (12 trials; including 2 sets explicit reports for familiarization). The instructions included, through the backstory, complete information of the task structure to minimize effects of structure learning during the early trials / blocks. Specifically, subjects were instructed of (a) the 3 reward levels, volatility (changes every 24 trials on average), and the fact that reward levels would change independently across optionarms and independently of the subjects’ responses, without accompanying cues (b) the noisiness of the obtained rewards about the mean reward levels, as well as the two levels of noise, (c) the presence of interleaved forced choice trials, and (d) the occasional explicit reports. Subjects were also instructed that their goal is to maximize rewards (total number of points obtained), although their actual compensation was not dependent on task performance. Following the full-block practice, subjects were asked a series of questions to ensure task comprehension, and verbal clarification was provided as needed.

In the second session, prior to going inside the MRI scanner, subjects completed another short practice block (12 trials) on a laptop computer, as a reminder of the task structure, as well as to familiarize them with the trial timing differences from the behavioral session, and the different response mappings in the scanner (one-hand in the scanner vs. both hands in the behavioral session).

### 2.3. Psychometric scales

Between the two sessions, at a time of their choosing, subjects completed an online battery of psychometric scales comprising previously validated French translations of the following 8 questionnaires:

- Weiss Functional Impairment Rating Scale (WFRIS) for Attention Deficit and Hyperactivity Disorder (ADHD), with subscales for different domains including family, work, school, life skills, self-concept, social, and risk-taking ^76^; French translation ^77^
- State-trait anxiety inventory, form Y (STAI-Y) for clinical anxiety, with subscales for state and trait anxiety ^68^; French translation ^78^
- Revised Life Orientation Test (LOT-R) for optimism ^79^; French translation ^80^;
- Barratt Impulsiveness Scale (BIS-11) for impulsivity ^81^; French translation ^82^;
- Snaith-Hamilton Pleasure Scale (SHAPS) for anhedonia ^83^; French translation ^84^;
- Autism quotient (AQ) for autism spectrum ^85^; French translation ^86^
- Schizotypal personality questionnaire (SPQ) for schizophrenic symptomatology ^87^; French translation ^88^
- Big Five Personality Inventory (BIG 5), measuring the “Big 5” personality traits: openness, conscientiousness, extraversion, agreeableness, and neuroticism ^89^; French translation ^90^.

The scales were presented in a fixed order, as listed above, across all subjects, on the Gorilla Experiment Platform (https://gorilla.sc/). The French translations of the questionnaires are available on github <add link>.

### 2.4. Models

#### Learning model: Bayesian ideal observer

The Bayesian learner was specified in a Hidden Markov Model (HMM) framework. Starting from a uniform prior over the 3 reward levels for each option p(μ_0_), on each trial, the model updates p(μ^(k)^_i,t+1_|y_1:t+1_, σ_1:t+1_, ν), the posterior probability of the hidden reward level μ_i_ corresponding to option *k* using the new outcome y_t+1_, and assuming some fixed probability ν of a changepoint, specified via a transition matrix (with 1-p(changepoint) on the diagonal; p(changepoint) evenly distributed over the remaining columns of each row). The likelihood p(y_t+1_|μ^(k)^_i,t+1_, σ_t+1_) evaluates the probability of the current outcome y_t+1_ following the choice of option *k*, at each reward level μ_i_, using a normal density function (with a mean μ_i_ and the standard deviation σ_t+1_ corresponding to the generative noise level. Note that when no outcome is available for a given option, which occurs whenever this option was not selected, the likelihood function is constant (it does not depend on μ_i, t+1_). The update relies on Bayes rule and leverages the fact that, conditioned on the previous reward level μ^(k)^_i,t_, the new reward level μ^(k)^_i,t+1_ is independent from previous outcomes y_1:t_, and the fact that, conditioned on the current reward level μ_i,t+1_, the current outcome y_t+1_ is independent from the previous ones y_1:t_. The update equation for the posterior is:

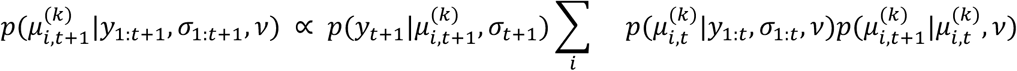

We used this posterior to model the maximum a posteriori reward level and estimation uncertainty at the moment of questions.

For choices, we modeled the expected reward level, estimation uncertainty and prediction error using the predictive distribution of reward levels (by updating the posterior from one trial to the next assuming that a change point could occur):

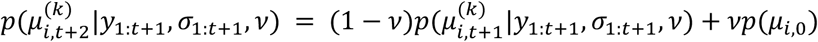

#### Learning model validation

A key reason for using a Bayesian learner in this study is that it provides trial-by-trial estimates of uncertainty, which can be used to assess subjects’ uncertainty-driven behavior. To validate that this learning model adequately fits the reward estimates of subjects, we compared the Bayesian learning model against a Rescorla-Wagner (RW) delta-rule learning model ^47,91^. The standard delta-rule learning model updates the reward estimate of the chosen option only. Given the poor performance of this model, we used a variant to also update the value of the unchosen option, making it decay toward the average reward level (50):

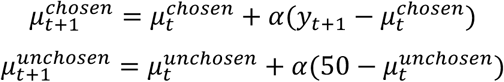

Where α is the learning (and forgetting) rate, fit to the subjects choice (assuming as a choice model the baseline model presented in the Result section).

#### Model of choice stochasticity

We considered that choices (repeat or switch) are a stochastic function of a linear combination of decision factors *f_i_*. We distinguished two forms of stochasticity. One assumed that choice probability increases following the linear combination of decision factors. More specifically, choice probability is a logistic function of the linear combination of factors (when there are two possible choices, this choice model is related to the softmax policy^47^):

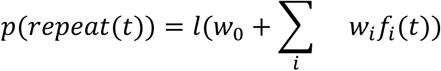

With *l* the logistic function: 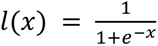

The other model of choice stochasticity, known as epsilon-greedy^47^ assumed that subjects chose to repeat if *w*_0_ + ∑*_i_ w_i_f_i_*(*t*) is positive (and to switch otherwise) with probability 1-ε (constant and independent across trials) or else chose the other option.

#### Decision factors

The decision factors were derived from the ideal observer posterior probabilities of the reward levels:

- *Relative expected reward (ΔER)*. It is the difference (repeat minus stay) in expected reward associated with each option. The expected reward for option *k* on trial *t* is computed from the predictive distribution of reward levels: 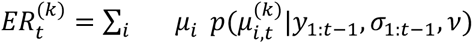.
- *Estimation uncertainty (ΔEU* and *EUt)*. We modeled the estimation uncertainty of the reward levels of option *k* on trial *t* as the residual probability mass that is not the maximum predictive probability across reward levels: 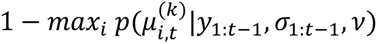. This quantity is particularly sensitive to the noise level and recent changepoints, as well as the option sampling history. Two aspects of estimation uncertainty were considered: relative (ΔEU; i.e., the difference between options) and total (EUt; i.e., the average across options; it captures a modulation of the repetition bias by total estimation uncertainty).
- *Signed and unsigned prediction error on the previous trial (PE and UPE)*. The prediction error is the difference between the expected and obtained reward on the previously chosen arm 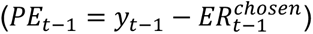. When controlling for the other factors the effects of the signed and unsigned predictor error reflect the second heuristic of interest in the study: a “win-stay-lose-shift” strategy, whereby decisions on the current trial are based on the direction and amount of deviation of the outcome of the previous choice from expectation. The effect of the unsigned prediction error (|PE_t-1_|) captures a potential asymmetry in the slope of the PE effect between the positive and negative domains (see Results and Fig. S8).

We considered a baseline model consisting of only the constant w_0_ and *ΔER*. We then considered more complex models by adding each factor and each combination of factors in a full factorial way.

Therefore, the baseline model was:

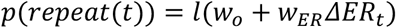

And the most complex main-effects model was:

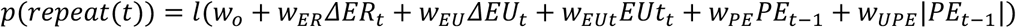

#### Interactions

To examine potential interactions of uncertainty (ΔEU, EUt) with other factors (ΔER, PE, UPE), combinations of candidate two-way interactions were added as additional regressors to the winning main-effects-only model, and the best-fitting model was the final selected model, on which the main results focus.

#### Model fit

Regressors were standardized (z-scored) prior to model fitting for comparability of the resulting coefficient estimates. Model fitting was performed in Python, with scipy.optimize, using maximum-likelihood estimation (MLE) on the log-loss between model predictions and subjects’ choices. For models that fitted volatility on top of the decision parameters, the Powell optimization procedure was used. For models that used the generative volatility, only the decision parameters remained to be fitted; the decision model being equivalent to a logistic regression model, we fitted a logistic regression model using scipy’s defaults (the Broyden-Fletcher-Goldfarb-Shanno (BFGS) procedure, called using the wrapper statsmodels.logit was used as a backend).

#### Model comparison

Models were compared with leave-one-block-out cross-validation (i.e., 11 blocks training, 1 block testing). More precisely, we used the cross-validated average choice likelihood, that is, the average model-derived probability of subjects’ choices on each free trial. The average choice likelihood is a sensitive measure of model fit for comparison, because it is less impacted by occasional low-probably choices than the quantity being minimized, and it has an intuitive interpretation: it reflects the extent to which subjects’ choices are, on average, consistent with model predictions. For the main goal of comparing decision models with different subsets of candidate factors, cross-validation was performed on the decision-model only, using subject-specific volatility estimates derived on all data (rather than fitting on the training set only). This was done to reduce the amount of computing time needed to fit the models. As an additional check, after the final main-effects model was selected, single-factor ablations were performed from this final model with full cross-validation (including the fit of volatility on the training set only). The final model reliably outperformed all ablated models except for a marginal contribution of one factor, the total estimation uncertainty (Δp(choice) = 0.001, *t*(55) = 1.92, *p* = 0.06, Cohen’s *d* = 0.26) (see Fig. S12 for full results).

#### Parameter and model recovery

Parameter recovery was performed to ensure that effects in the selected model are sufficiently dissociable to estimate independent coefficients. The parameter recovery procedure was as follows: (1) sample parameter values uniformly within the range of subjects’ actually observed parameter estimates, independently for each parameter; (2) simulate choices with these parameter settings; (3) fit a model to the simulated choices to obtain recovered parameter values; (4) iterate 1000 times; and (5) compute Spearman correlations of the generative and recovered parameter estimates across iterations, for all pairs of parameters, to produce a confusion matrix. (Fig. S1)

For the model recovery procedure, steps (1)-(2) were identical. For (3), all main-factor models (n=16) were fit to choices generated with a given model; for (4), there were 100 iterations. To evaluate the results, (5), we counted the number of times a given generative model was best fit by each model. (Fig. S2)

##### Statistical analysis

The group-level inferential statistics were as follows. We performed one-sample (against 0, for e.g., on logistic regression coefficients; or chance level, where different from 0; for e.g, on proportion of explicit reports matching the Bayesian learner) and paired-sample t-tests (for e.g., on cross-validated model fits for pairs of models). Reported correlation coefficients are either Pearson’s product-moment correlations or Spearman’s rank correlations, as noted in the text.

### 2.5. Code and data

Code and data are available here (see README.txt).

This copy of the code and data is for peer review purposes. Please do not share this link.

## Acknowledgements

This work was funded by CRCNS grant NIH/NIDA R01DA050373 to AY; ANR 19-NEUC-0002-01 to FM. We thank the nurses, radiographers and other staff at NeuroSpin, INSERM-CEA Cognitive Neuroimaging Unit for support with data collection.

## Contributions

FM, AY, DG, and AP conceived and designed the study; FM & AY supervised the project; ML & AP collected the data; FM & DG contributed models, AP analyzed the data; AP, ZH, AY, & FM prepared the study for publication. AP wrote the manuscript with discussion and comments by all co-authors.

## Supplementary figures

**Figure S1:**
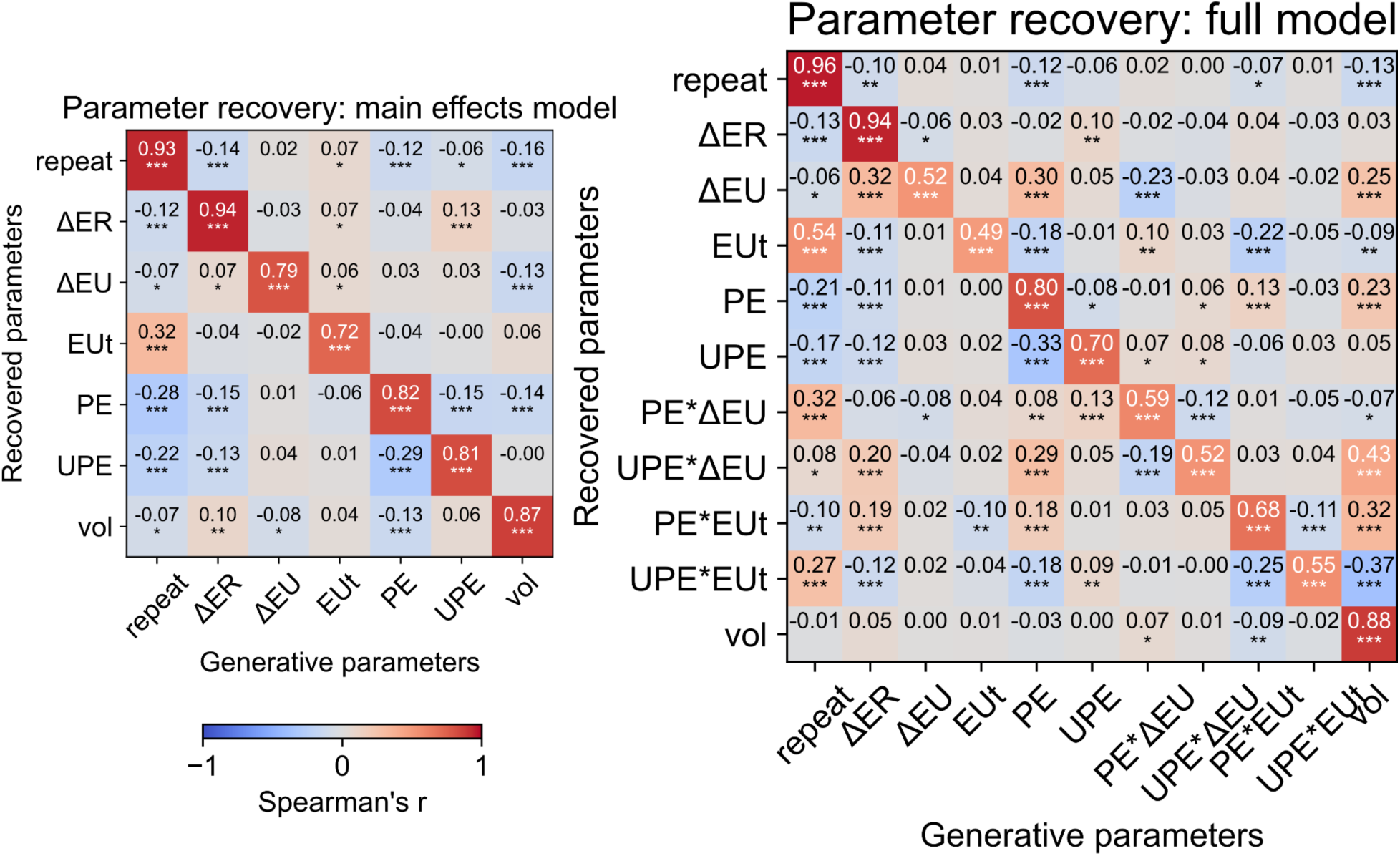
Parameter recovery. Spearman’s correlations between generative and recovered parameter estimates across 1000 iterations of simulated choices for the main effects model (left) and the full model, including interactions (right). *p < 0.05, **p < 0.01, ***p < 0.001.

**Figure S2.**
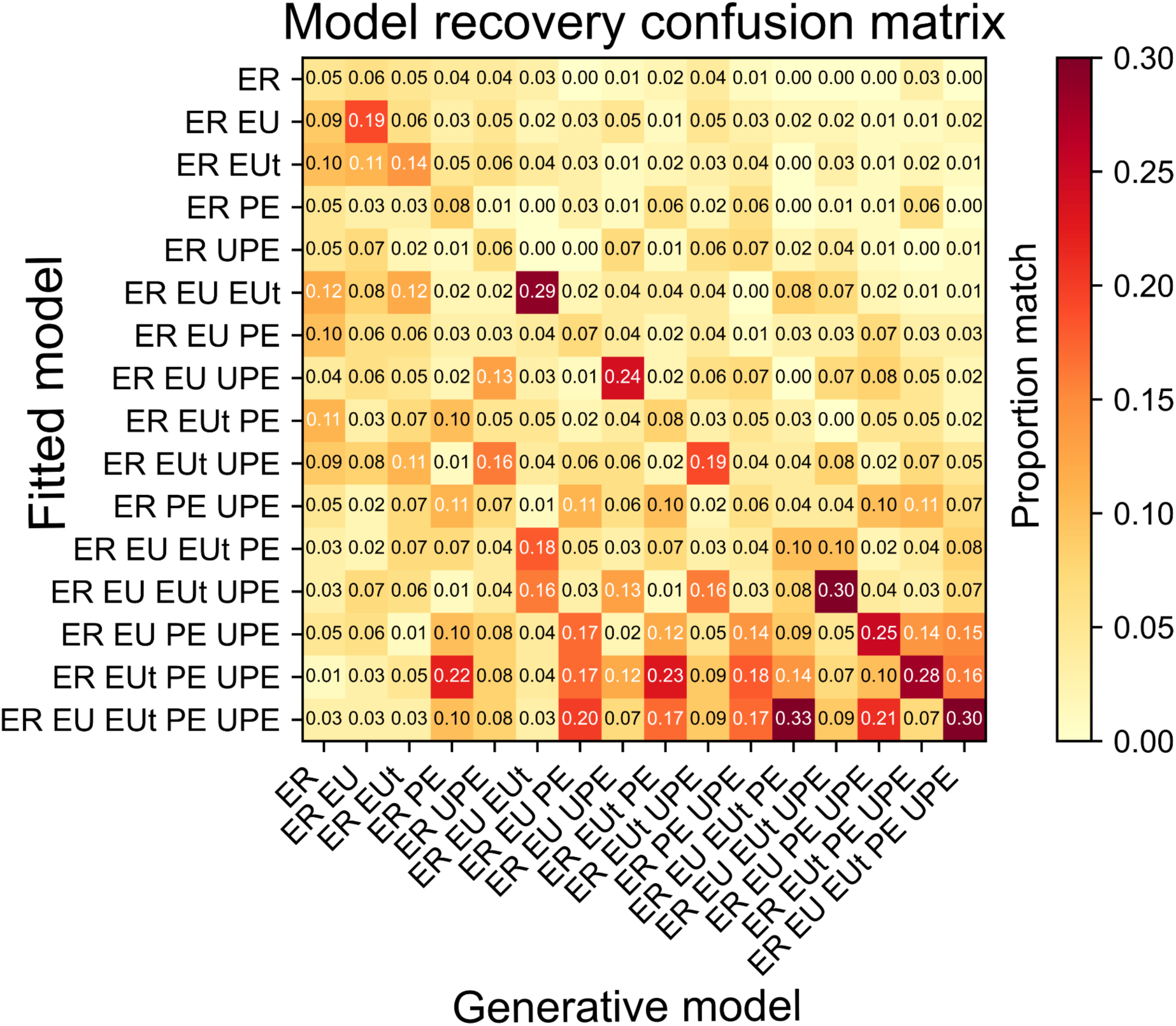
Model recovery. Confusion matrix of proportion of simulated choice sequences (out of 100 iterations) generated with a given model (x-axis) that are best fitted by each model (y-axis) (including only main effects models; n=16). Note that the larger proportion of matches in the lower-left triangle compared to the upper-right triangle of the matrix indicates that behavior generated by a given model can be well accounted for by a more complex model, but not by a simpler model. In particular, behavior generated by the most complex model with all main effects (which best accounted for our subjects’ data) is not well fitted by any of the simpler models.

**Figure S3.**
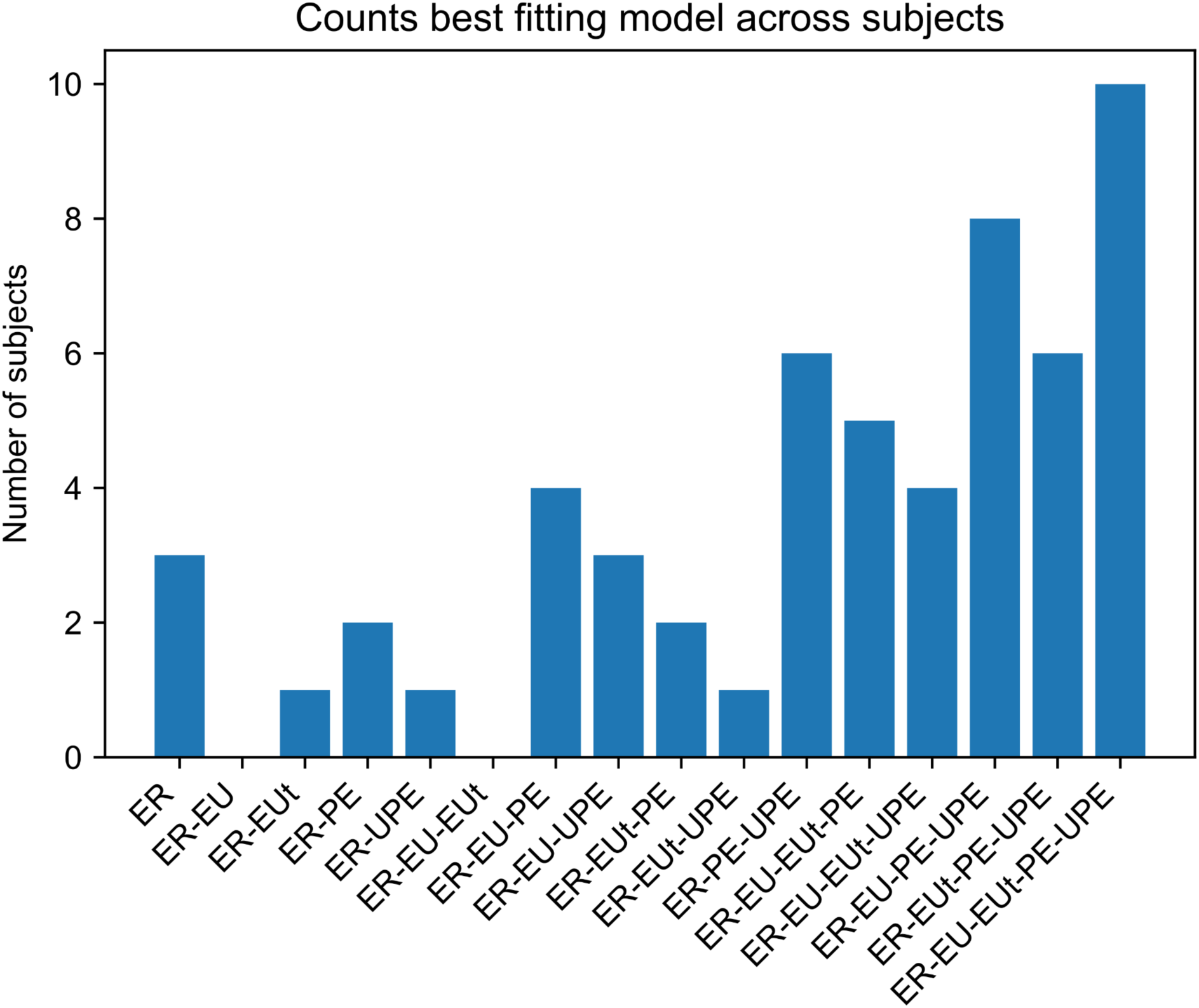
Counts of best fitting model across subjects. Number of subjects who are best fitted by each of the main effects models. Note that the full model and the models excluding only either ΔEU or EUt are the best-fitting models for 24 of 56 subjects (43%), and models that include one or more uncertainty term best-fitted 44 of 56 subjects (79%).

**Figure S4.**
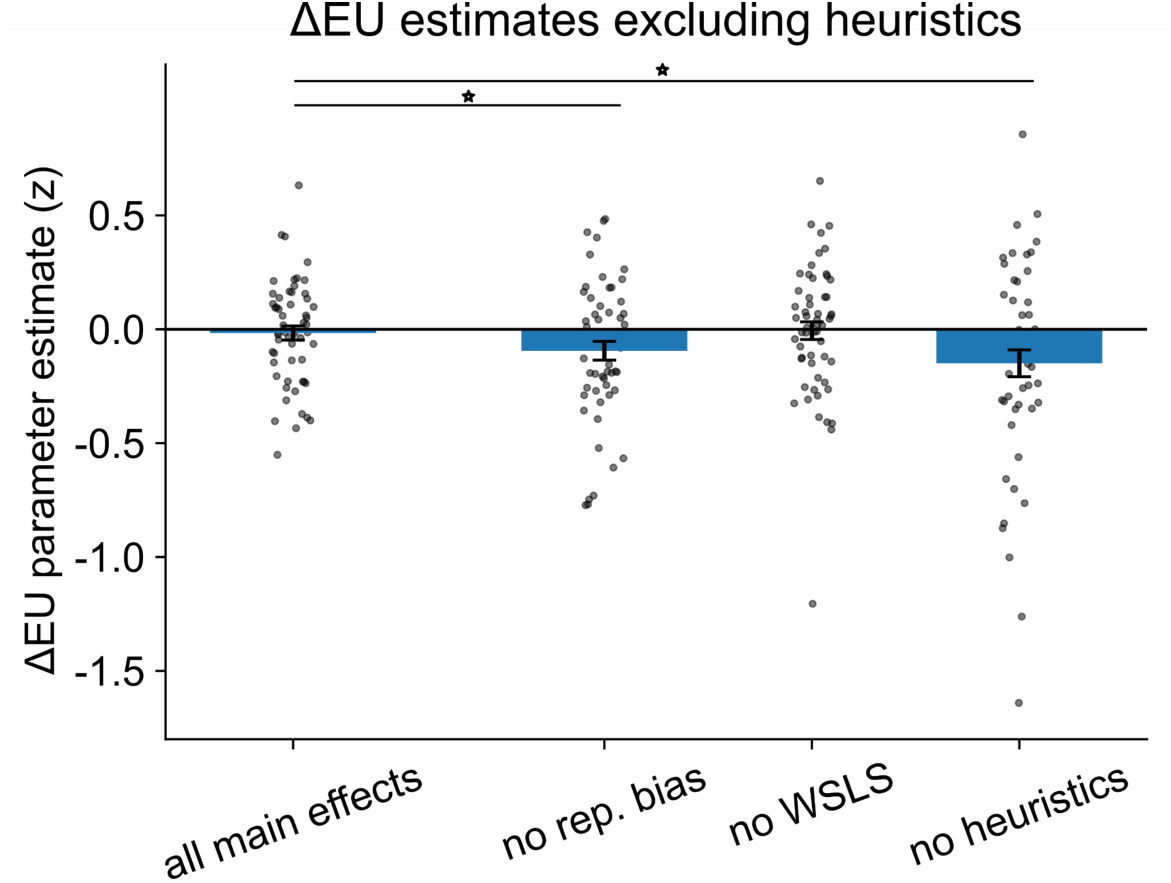
ΔEU parameter misestimation without heuristics. Omitting either the repetition bias (“no rep. bias”) or both repetition bias and win-stay-lose-shift heuristic terms (PE and UPE) (“no heuristics”) results in significantly different group-level ΔEU effects (see also Fig. S5). This suggests ΔEU can be misestimated if the heuristics are not accounted for.

**Figure S5.**
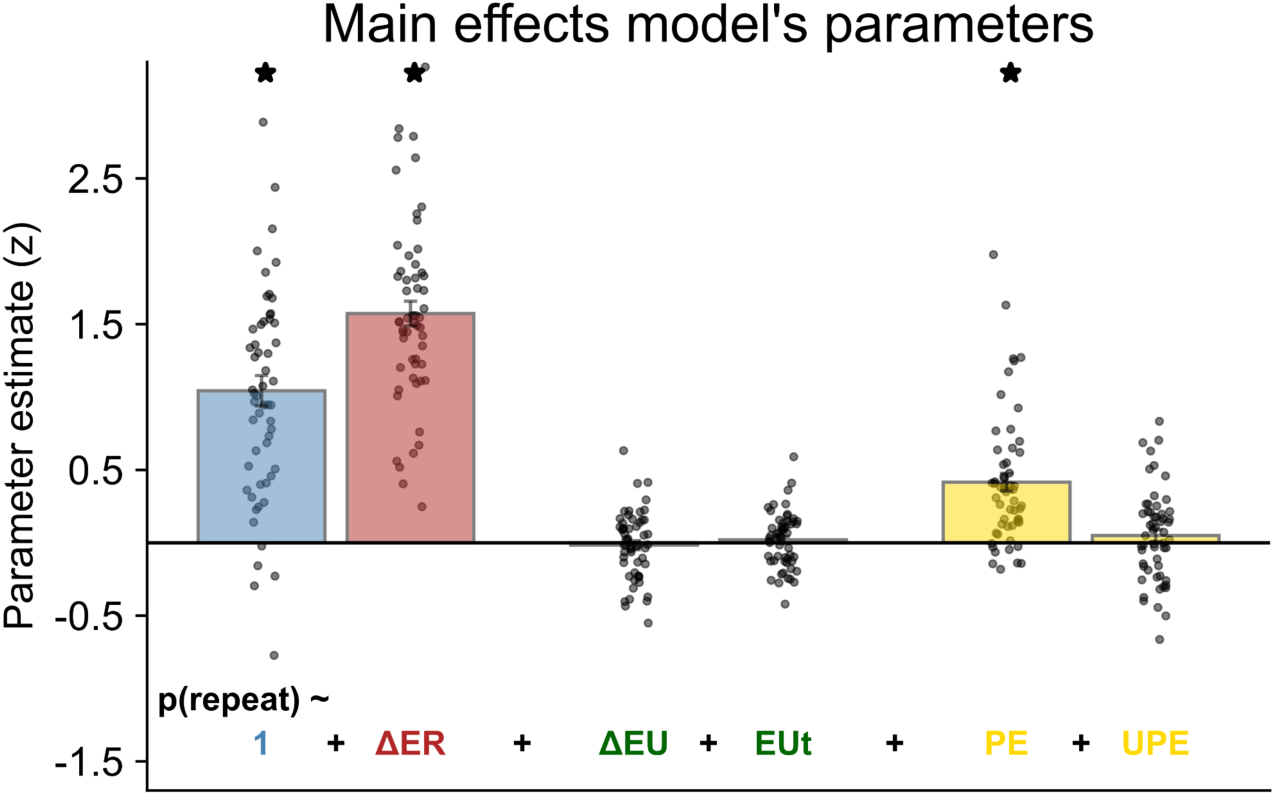
Main effects model. Parameter estimates for the model with main effects only (no interactions). * indicates p < 0.05.

**Figure S6.**
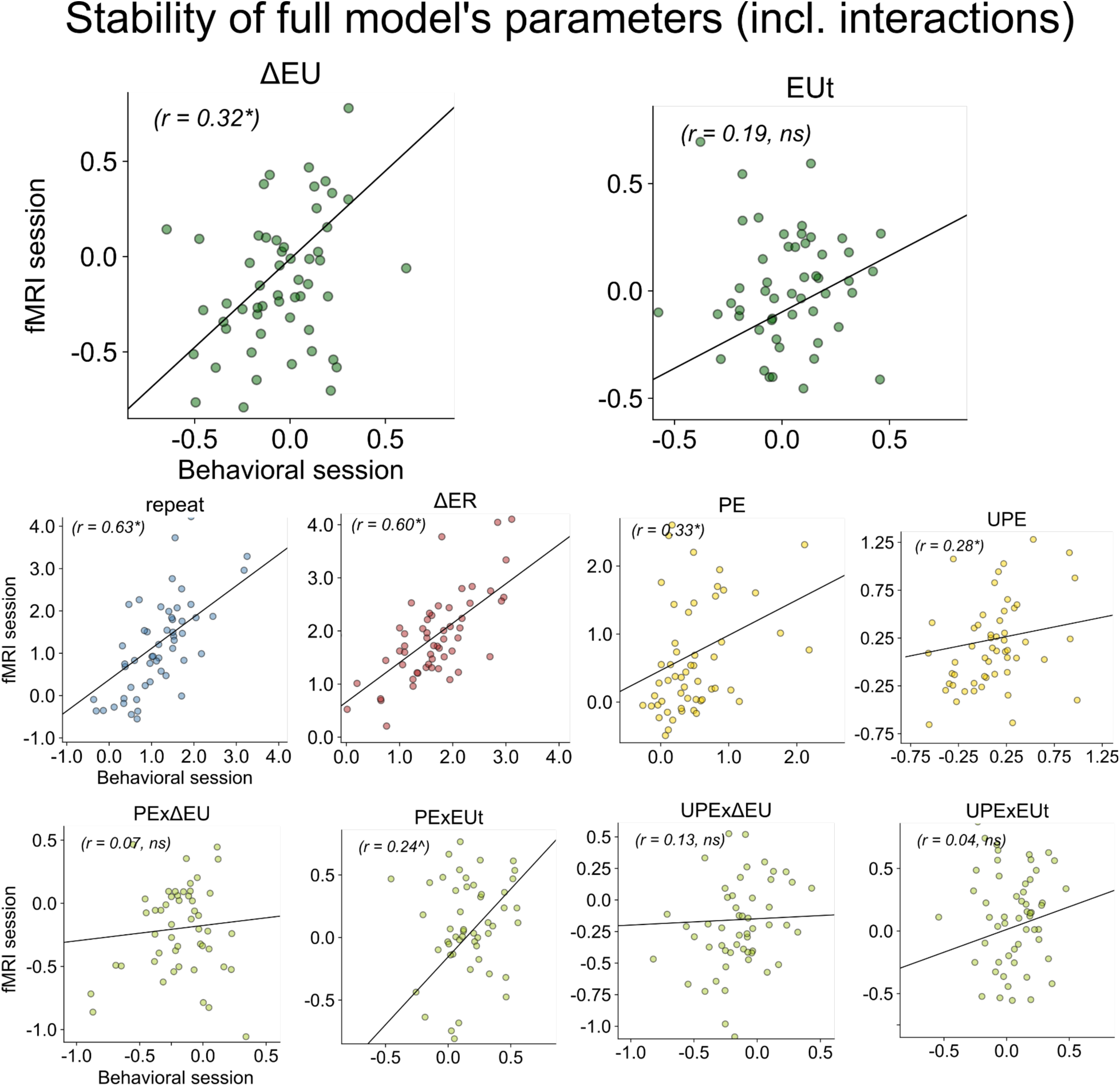
Parameter stability of the model including interactions. Spearman’s rank correlations of parameter estimates across testing sessions for full model, including interactions. *p < 0.05; ^p < 0.1.

**Figure S7:**
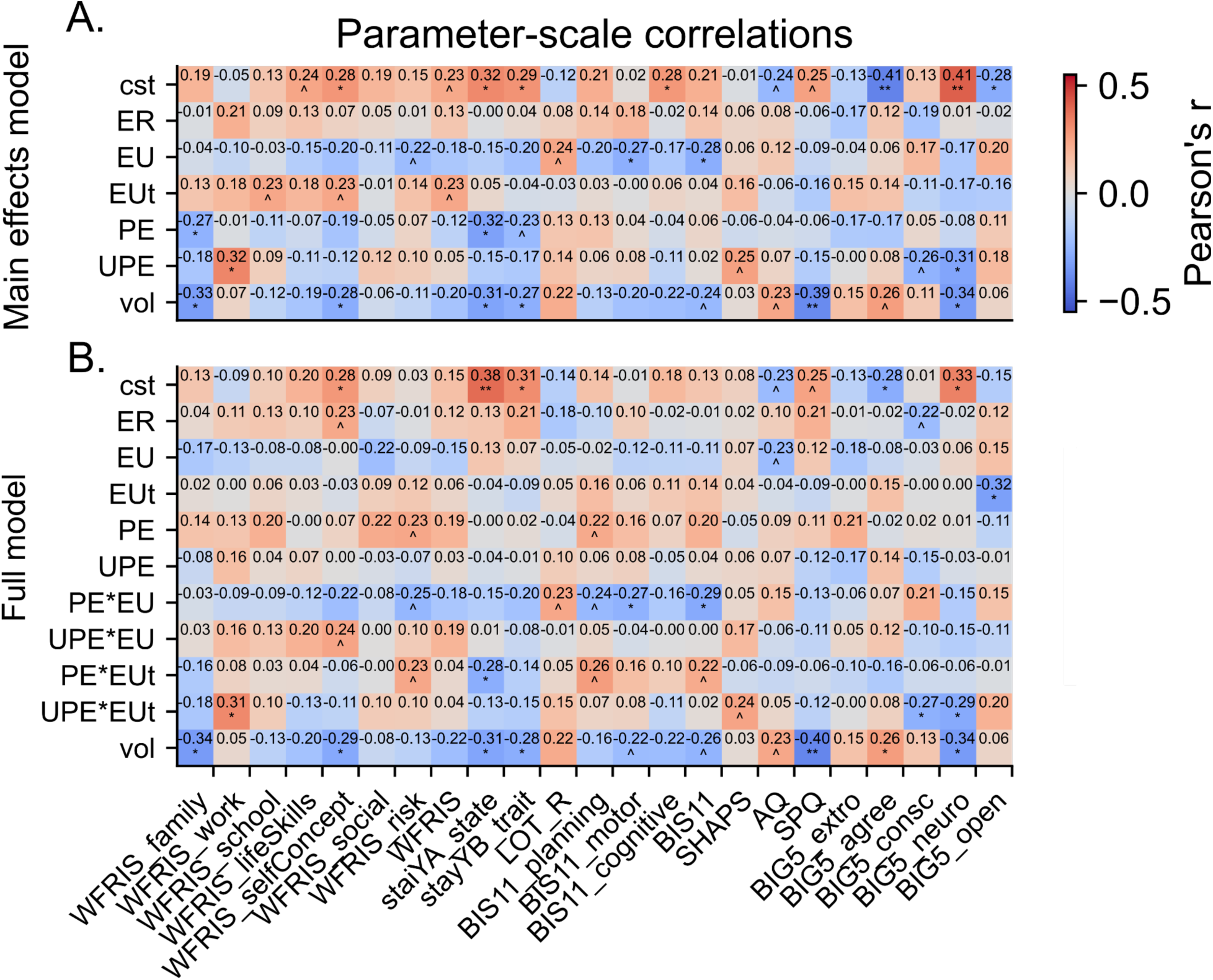
Psychometric scales correlations with parameter estimates from the models. Pearson correlations of all included psychometric scales and subscales with **A:** parameters of main effects-only model and **B:** parameters of full model including interactions. ^p < 0.10; *p < 0.05, **p < 0.01. Scale acronyms: *WFRIS*: Weiss Functional Impairment Rating Scale; *STAI-Y*: State-Trait Anxiety Inventory; *LOT-R*: Life Orientation Test - Revised; *BIS-11*: Barratt Impulsiveness scale. SHAPS: Snaith–Hamilton Pleasure Scale; AQ: Autism quotient; SPQ: Schizotypal personality questionnaire; BIG 5: Big five personality questionnaire (extroversion, agreeableness, conscientiousness, neuroticism, openness to experience).

**Figure S8.**
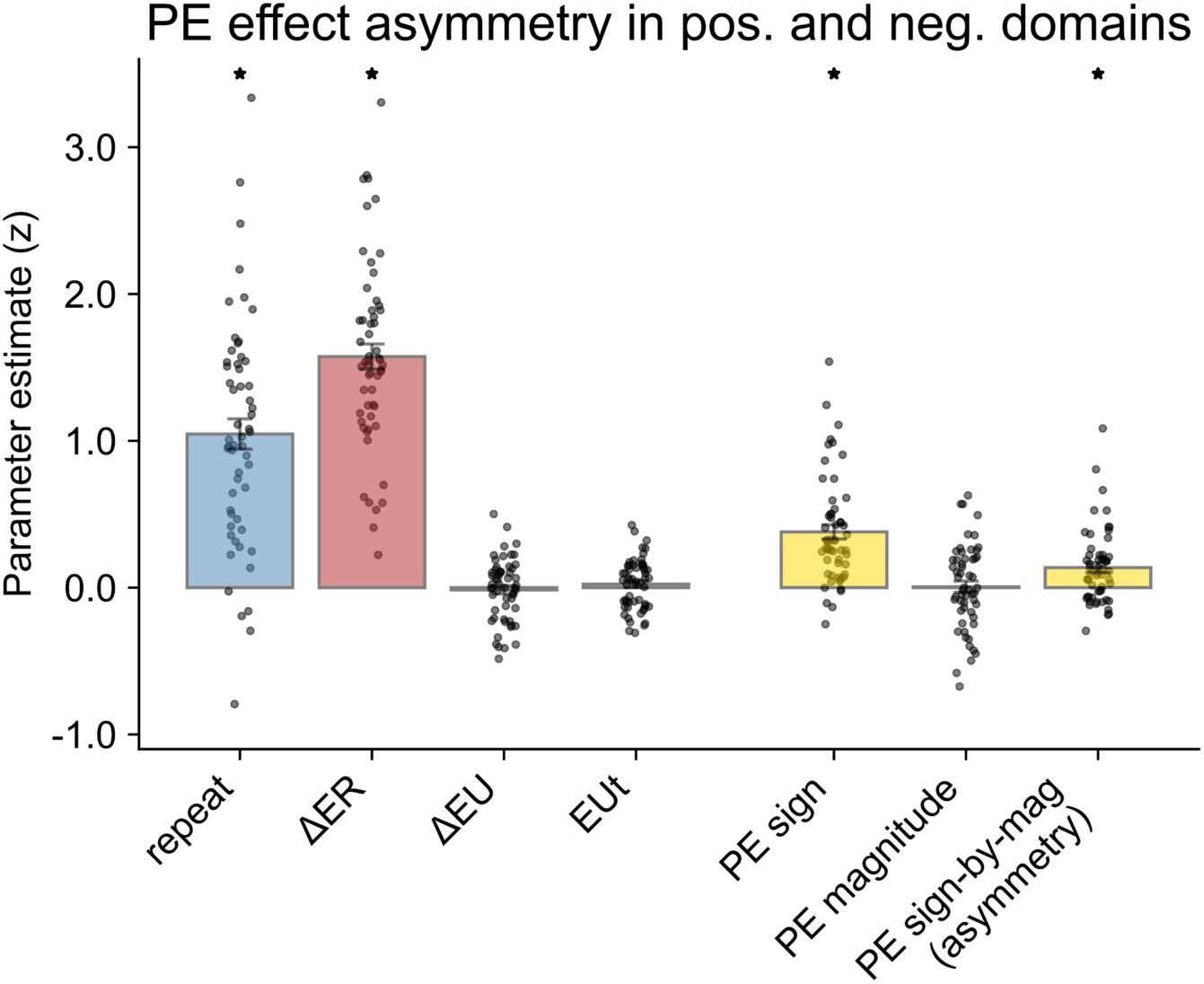
Asymmetry of prediction error effect. Test of the interpretation of the unsigned prediction error term (UPE) as capturing an asymmetry effect, as used in the model presented in the main text. The previous prediction error (PE, capturing win-stay-lose-shift heuristic) in the main-effects model is parameterized into “sign” (indicator 0/1 variable), “magnitude” (i.e., unsigned prediction error), and interaction terms (sign x magnitude). The interaction term captures the asymmetry of the PE effect in the positive and negative domains (beta = 0.137, SE = 0.034, *t*(55) = 4.037, *p* = 0.0002, *Cohen’s d* = 0.54). *p < 0.05. In the main text, this asymmetry is captured by the UPE (unsigned prediction error). The advantage of a model with PE and UPE over PE sign, PE magnitude and their interactions is model simplicity and easier modeling of interaction with uncertainty.

**Figure S9.**
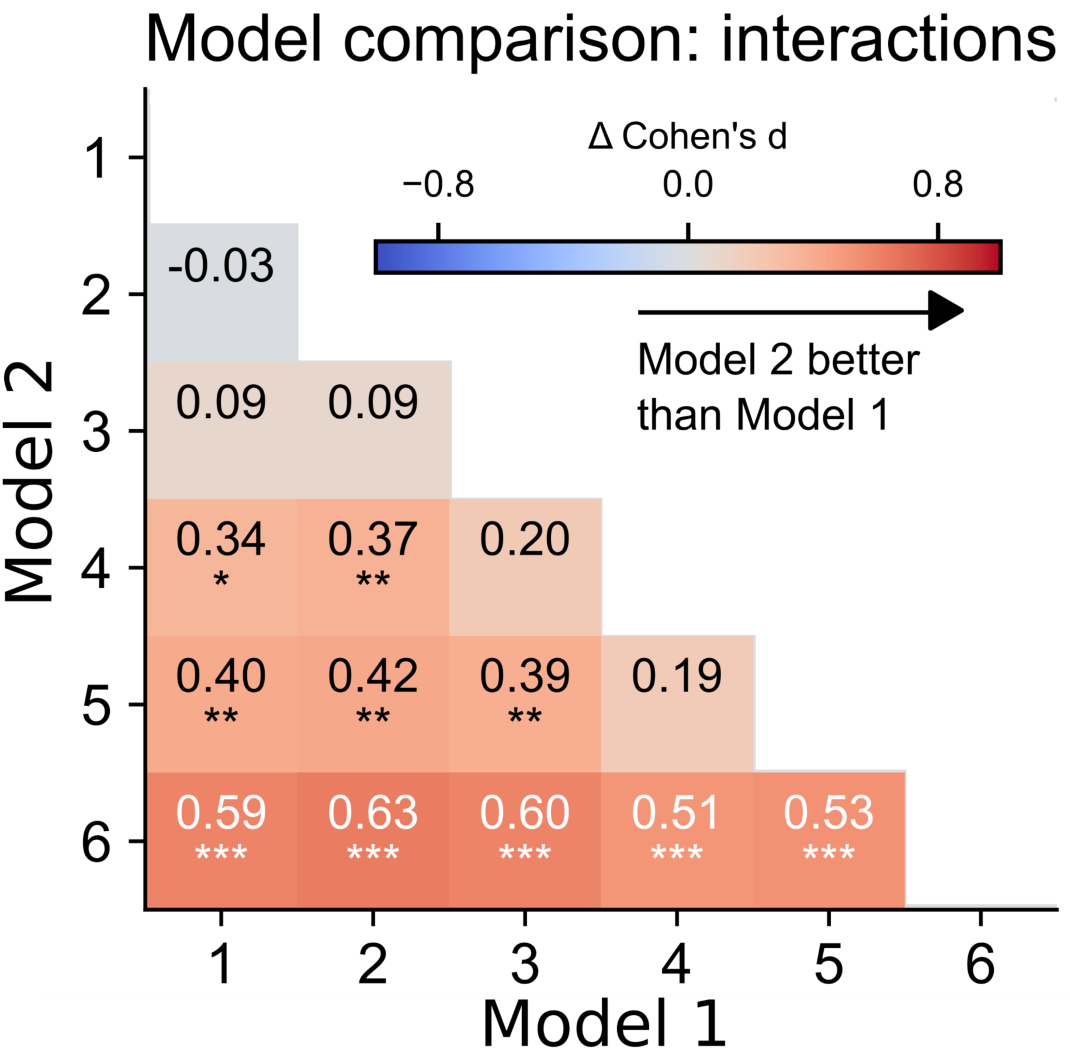
Model selection with interactions. Model fit improvements from adding interaction terms to the selected main effects model. The mode complex model is better than any simpler model. Model numbers correspond to: 1. repeat ∼ 1+ΔER+ΔEU+EUt+PE+UPE (base model without interactions) 2. repeat ∼ 1+ΔER+ΔEU+EUt+PE+UPE**+ΔER*ΔEU** 3. repeat ∼ 1+ΔER+ΔEU+EUt+PE+UPE**+ΔER*EUt** 4. repeat ∼ 1+ΔER+ΔEU+EUt+PE+UPE**+PE*ΔEU+UPE*ΔEU** 5. repeat ∼ 1+ΔER+ΔEU+EUt+PE+UPE**+PE*EUt+UPE*EUt** 6. repeat ∼ 1+ΔER+ΔEU+EUt+PE+UPE**+PE*ΔEU+PE*EUt+UPE*ΔEU+UPE*EUt** *p < 0.05, **p < 0.01, ***p < 0.001.

**Figure S10.**
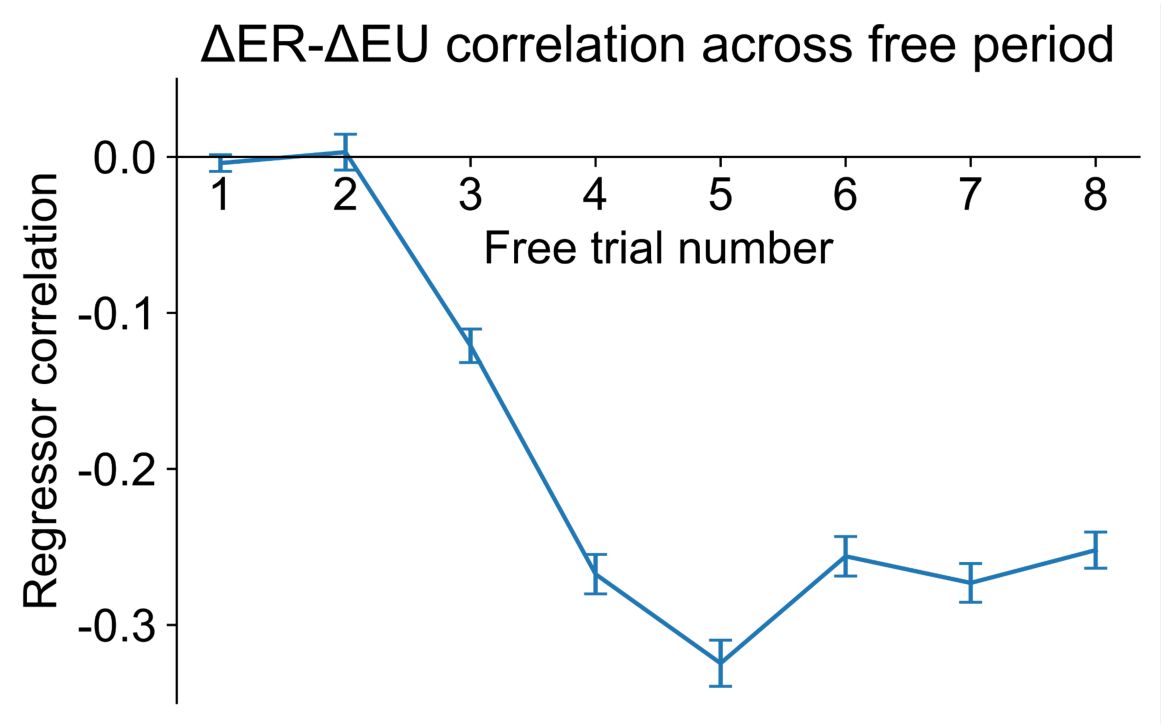
Decorrelation of reward and uncertainty by forced choices. Anticorrelation (Pearson’s *r*) of ΔER and ΔEU on each free trial following a forced choice period. This analysis reveals the benefit of interleaving periods of free and forced choices: a negative correlation between ΔER and ΔEU builds up within periods of free trials, which forced choices (temporarily) abolish.

**Figure S11.**
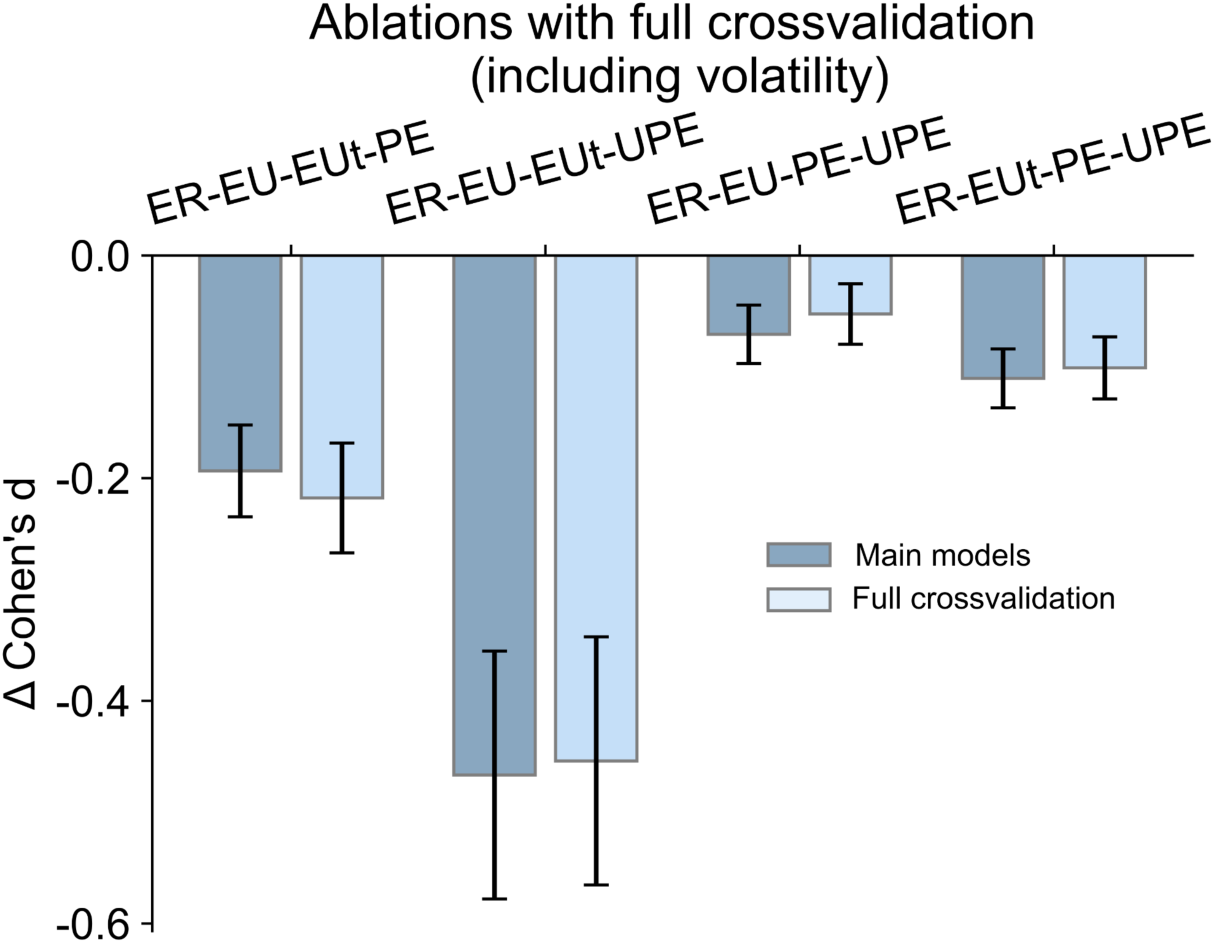
Ablation tests with full vs partial cross-validation. Decrements in model fit relative to model with all main effects (*repeat ∼ 1+ΔER+ΔEU+EUt+PE+UPE*), from removing each additional factor (UPE, PE, EUt, ΔEU, left to right), for “Main models”, reported in the text (excluding volatility) and with “Full crossvalidation” (including volatility parameter). This analysis shows that similar conclusions can be made when fitting all parameters (including volatility) compared to the simpler version adopted in the main text, where volatility is not fitted in each fold of the crossvalidation.

**Figure S12.**
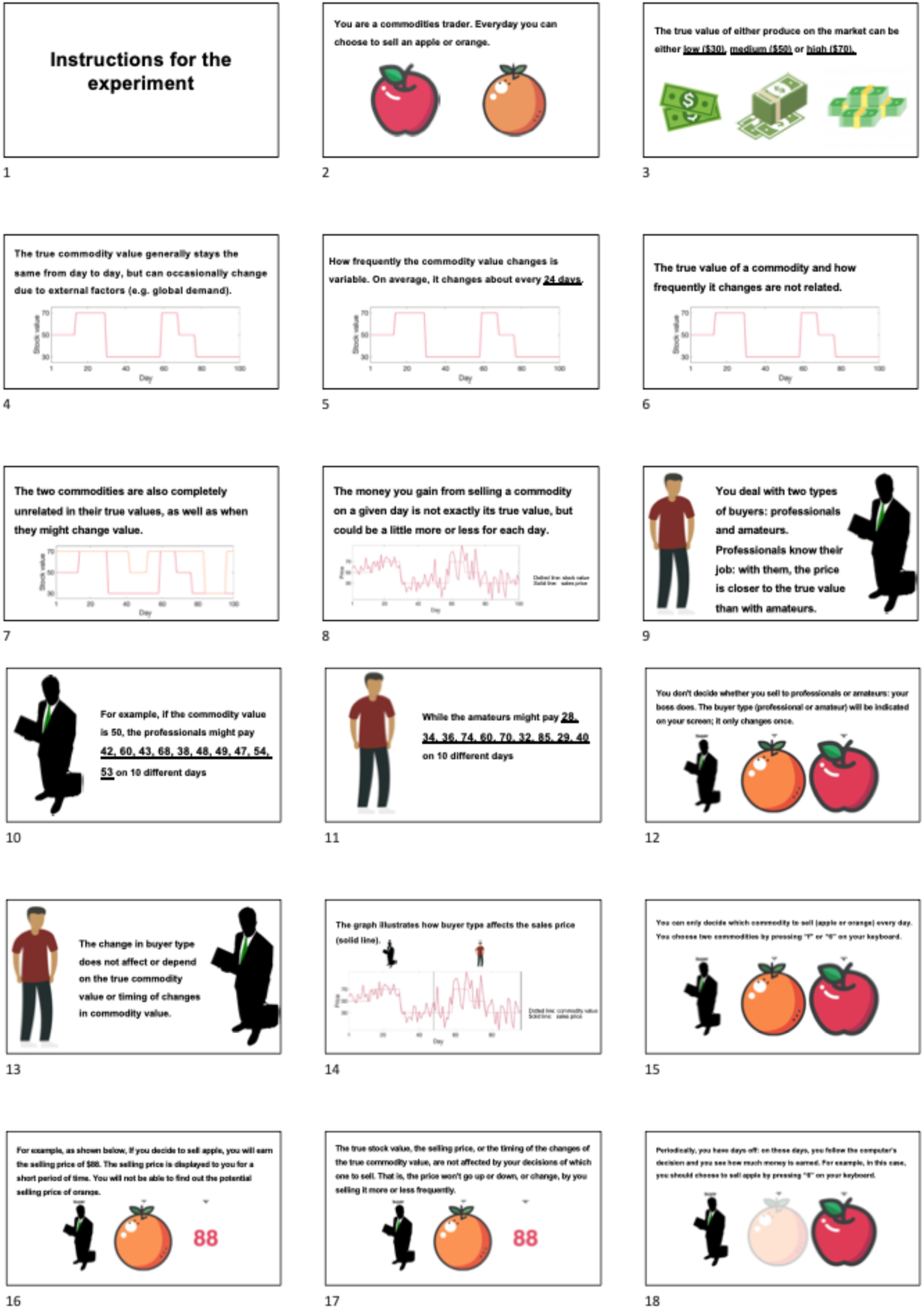
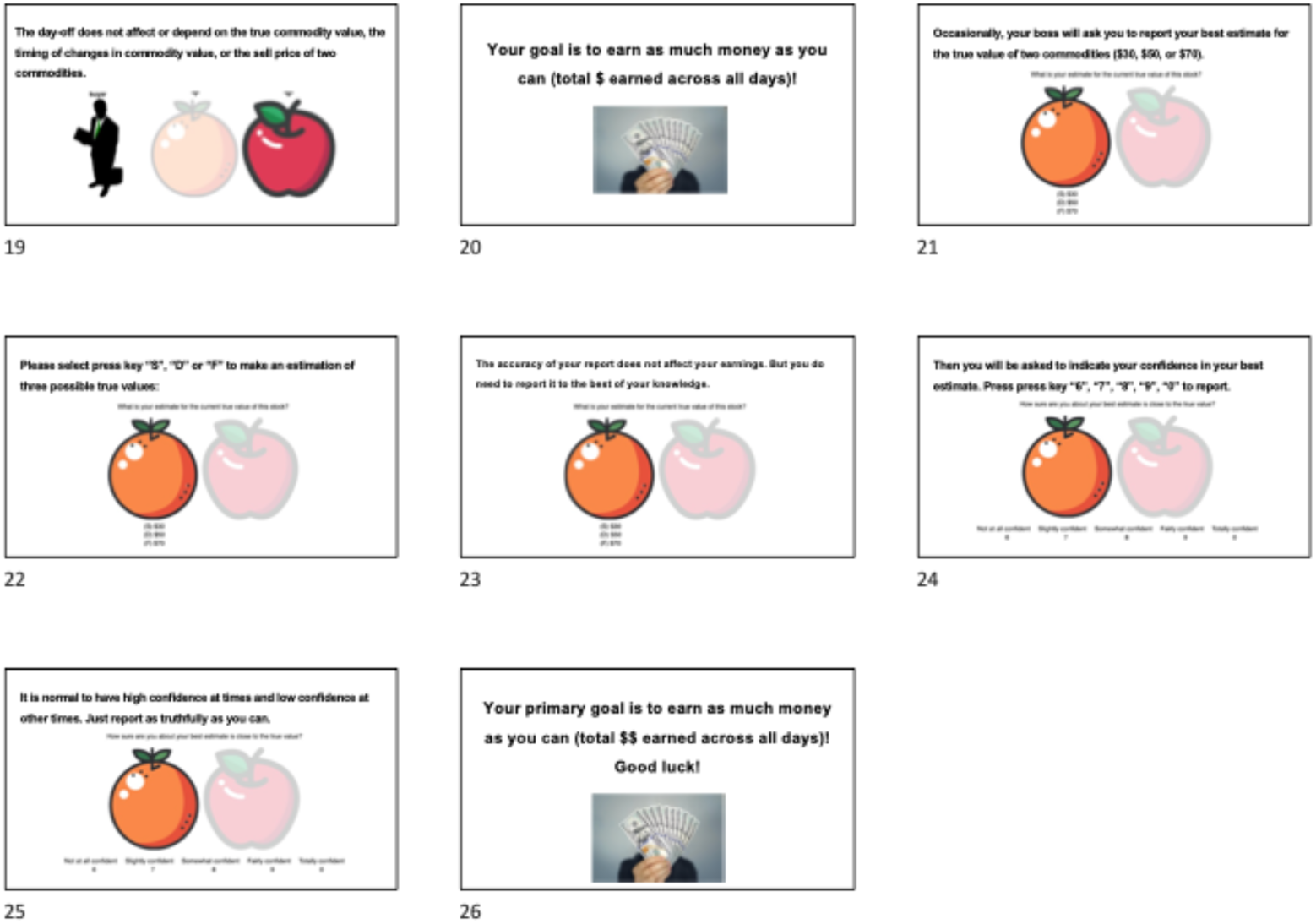
Instructions for illustrated version of the task with back story (translated from French to English).

